# Neuronal GPCR regulates the specificity of innate immunity against pathogen infection

**DOI:** 10.1101/2021.05.06.442992

**Authors:** Phillip Wibisono, Shawndra Wibisono, Jan Watteyne, Chia-Hui Chen, Durai Sellegounder, Isabel Beets, Yiyong Liu, Jingru Sun

## Abstract

A key question in current immunology is how the innate immune system generates high levels of specificity. Using the *Caenorhabditis elegans* model system, we demonstrate that functional loss of NMUR-1, a neuronal G protein-coupled receptor homologous to mammalian receptors for the neuropeptide neuromedin U, has diverse effects on *C. elegans* innate immunity against various bacterial pathogens. Transcriptomic analyses and functional assays revealed that NMUR-1 modulates *C. elegans* transcription activity by regulating the expression of transcription factors involved in binding to RNA polymerase II regulatory regions, which, in turn, controls the expression of distinct immune genes in response to different pathogens. These results uncovered a molecular basis for the specificity of *C. elegans* innate immunity. Given the evolutionary conservation of NMUR-1 signaling in immune regulation across multicellular organisms, our study could provide mechanistic insights into understanding the specificity of innate immunity in other animals, including mammals.

## Introduction

The pathogen-triggered host immune response is a multilayered process programed to fight invading microorganisms and maintain healthy homeostasis. There are two types of immune systems: the innate immune system and the adaptive immune system. The innate immune system, comprised of physical barriers as well as inducible physiological and behavioral responses, serves as the first line of defense for the host and is evolutionarily conserved across multicellular organisms. Innate immune responses are initiated by recognizing pathogen-associated molecular patterns (PAMPs) or damage caused by pathogen-encoded virulence factors (Jones and Dangl, 2006; Medzhitov, 2009). Such recognition triggers multiple internal defense responses, including the release of cytokines and chemokines, to fight invading microbes directly or indirectly. In vertebrates, when the innate immune response is insufficient to control an infection, the highly specific adaptive immune response is activated as the second line of defense (Chaplin, 2010). In this system, antigen-presenting cells (APCs) engulf and break down pathogens and present the foreign antigens on their cell surface via MHC class II molecules. Subsequently, T cells and B cells with antigen-specific receptors are activated to destroy infected cells or produce antibodies to inhibit infection, respectively.

While a large repertoire of T- and B-cell receptors and antibodies in the adaptive immune system confer remarkable specificity against different pathogens, the innate immune response was traditionally thought to be non-specific. However, increasing evidence indicates that the innate immune system can also generate high levels of specificity. In vertebrates, a diverse array of pattern recognition receptors (PRRs), such as the Toll-like receptors (TLRs), NOD-like receptors (NLRs), retinoic acid-inducible gene-I-like receptors (RLRs), and C-type lectins (CLRs), recognize various PAMPs and mediate a broad pattern of specificity (Pees et al., 2016). The diversity of ligand binding can be further expanded by the formation of receptor homodimers or heterodimers through interactions with other PRR co-receptors (Tan et al., 2014). In addition to controlling immune activation through specific receptor-ligand binding, innate immunity is also tightly regulated by the nervous system to ensure that appropriate responses are mounted against specific pathogens. Such regulation is critical because insufficient immune responses can exacerbate infection and excessive immune responses can lead to prolonged inflammation, tissue damage, or even death (Steinman, 2004; Sternberg, 2006). The mechanisms underlying neuro-immune regulation, however, are not well understood and are difficult to decipher due to the high degree of complexity of the nervous and immune systems in most model organisms. Overall, the molecular basis responsible for the specificity of innate immunity in vertebrates remains unclear and is largely understudied, likely because such specificity is overshadowed by the concomitant adaptive immune response.

Unlike vertebrates, most invertebrates do not have an adaptive immune system and rely solely on innate immunity to defend themselves against pathogen infection, yet they can still differentiate between different pathogens (Pees *et al*., 2016). For example, *Drosophila melanogaster* possesses a repertoire of receptors that mediate the distinction of different types of pathogens and fungi (Cherry and Silverman, 2006). Alternative splicing of exons of the Dscam protein in insects generates thousands of isoforms that are potentially involved in pathogen recognition and resistance (Schulenburg et al., 2007). Moreover, in both *D. melanogaster* and *Caenorhabditis elegans*, brief pre-exposure of animals to a pathogen (termed priming or conditioning) provides strong protection against a subsequent challenge with an otherwise lethal dose of the same pathogen (Sharrock and Sun, 2020). These studies demonstrate that invertebrate innate immune systems can generate high levels of specificity, although the underlying mechanisms remain poorly understood. Nonetheless, the invertebrate systems allow for studies of the specificity of innate immunity without the confounding variability from adaptive immunity. In invertebrate organisms, the limited genetic diversity of PRRs and immune effectors is likely insufficient to explain the high levels of specificity (Schulenburg *et al*., 2007). This is exemplified by the case of *C. elegans*. The nematode genome does not encode the majority of known PRRs, and its sole TLR protein, TOL-1, does not seem to play key roles in immunity (Sun et al., 2016). However, *C. elegans* can still discern different pathogens and launch distinct immune responses against them (Irazoqui et al., 2010; Zarate-Potes et al., 2020). Different infections induce both pathogen-specific and shared immune genes in the worm (Wong et al., 2007; Zarate-Potes *et al*., 2020). While how the *C. elegans* immune system identifies specific pathogens remains elusive, several key studies have revealed that neural regulation of immunity has homologous occurrence in the nematode (Anyanful et al., 2009; Kawli and Tan, 2008; Pradel et al., 2007; Reddy et al., 2009; Styer et al., 2008; Sun et al., 2011; Zhang et al., 2005), which could control appropriate immune responses to different pathogens. The characteristics of *C. elegans*, such as the simplicity of its nervous and immune systems, genetic tractability, invariant lineage, effectiveness of RNA interference (RNAi), and a transparent body that allows for the monitoring of gene expression, make this organism uniquely suited for studying neural regulation of immunity. Indeed, such research in *C. elegans* is at the forefront of the field and has revealed unprecedented details regarding the molecules, cells, and signaling pathways involved in neural regulation of immunity (reviewed in (Liu and Sun, 2021)). For example, we have described an octopaminergic neuro-immune regulatory circuit in great detail (Liu et al., 2016; Sellegounder et al., 2018; Sun et al., 2012; Sun *et al*., 2011) as well as a neural-cuticle defense regulatory circuit mediated by the neuropeptide receptor NPR-8 (Sellegounder et al., 2019). Others have also revealed serotonergic, dopaminergic, cholinergic, and neuropeptidergic immunoregulatory pathways that control innate immune responses to various pathogens (reviewed in (Liu and Sun, 2021)). These studies have greatly improved our understanding of complex neuro-immune controlling mechanisms.

While the above-mentioned *C. elegans* studies showed how the nervous system regulates immunity against certain pathogens, they did not address whether or how such regulation generates specificity against different pathogens. During our work on neuro-immune regulation in *C. elegans*, we observed that functional loss of NMUR-1, a neuronal GPCR homologous to mammalian receptors for the neuropeptide neuromedin U (NMU), had diverse effects on the survival of *C. elegans* exposed to various bacterial pathogens. Furthermore, an earlier study by Alcedo and colleagues showed that an *nmur-1* null mutation affected the lifespan of *C. elegans* fed different *E. coli* food sources, and that effect was dependent upon the type of *E. coli* lipopolysaccharide (LPS) structures (Maier et al., 2010). These findings indicate that NMU/NMUR-1 signaling might mediate the specificity of immune responses to different pathogens. NMU is known to have roles in regulating immune responses in addition to its functions regulating smooth muscle contractions in the uterus, reducing food intake and weight, and modifying ion transport (Martinez and O’Driscoll, 2015; Ye et al., 2021). Indeed, three independent studies in mice demonstrated that NMU from enteric neurons directly activates type 2 innate lymphoid cells through NMUR1 to drive anti-parasitic immunity (Cardoso et al., 2017; Klose et al., 2017; Wallrapp et al., 2017). In the current study, we have discovered that NMUR-1 modulates *C. elegans* transcriptional activity by regulating the expression of transcription factors, which, in turn, controls the expression of distinct immune genes in response to different pathogens. Our study has uncovered a molecular basis for the specificity of *C. elegans* innate immunity that could potentially provide mechanistic insights into understanding the specificity of innate immune responses in vertebrates.

## Results

### Functional loss of NMUR-1 differentially affects *C. elegans* survival against various bacterial pathogens

Functional loss of NMUR-1 has distinct effects on the lifespan of *C. elegans* fed different *E. coli* food sources, and such effects are dependent upon the type of *E. coli* LPS structures (Maier *et al*., 2010). Given the toxic and immunogenic nature of LPS (Mazgaeen and Gurung, 2020), these LPS-dependent lifespan phenotypes suggest that NMUR-1 might mediate distinct immune responses to different pathogens. To test this notion, we examined the susceptibility of wild-type *N2* and *nmur-1* null animals (*nmur-1(ok1387)* strain) to six paradigmatic pathogens that included three Gram-negative bacteria (*Salmonella enterica* strain SL1344, *Pseudomonas aeruginosa* strain PA14, and *Yersinia pestis* strain KIM5) and three Gram-positive bacteria (*Enterococcus faecalis* strain OG1RF, *Microbacterium nematophilum* strain CBX102, and *Staphylococcus aureus* strain NCTC8325). Compared to wild-type animals, *nmur-1(ok1387)* animals exhibited enhanced survival against *S. enterica* and *Y. pestis*, reduced survival against *E. faecalis*, and showed no differences in survival against *P. aeruginosa*, *M. nematophilum*, and *S. aureus* (Fig. 1 and Fig. S1). In addition, with two commonly used worm food sources, namely, *E. coli* strains OP50 and HT115, we found that the *nmur-1* mutation improved worm survival against OP50 but had no effect against HT115 (Fig. 1 and Fig. S1), an observation in agreement with a previous study by Alcedo and colleagues (Maier *et al*., 2010). These distinct effects of the *nmur-1* mutation on survival, which appeared to be unrelated to the Gram-negative or Gram-positive groups, indicate that NMUR-1 plays diverse roles in *C. elegans* defense against various pathogens.

**Fig. 1.**
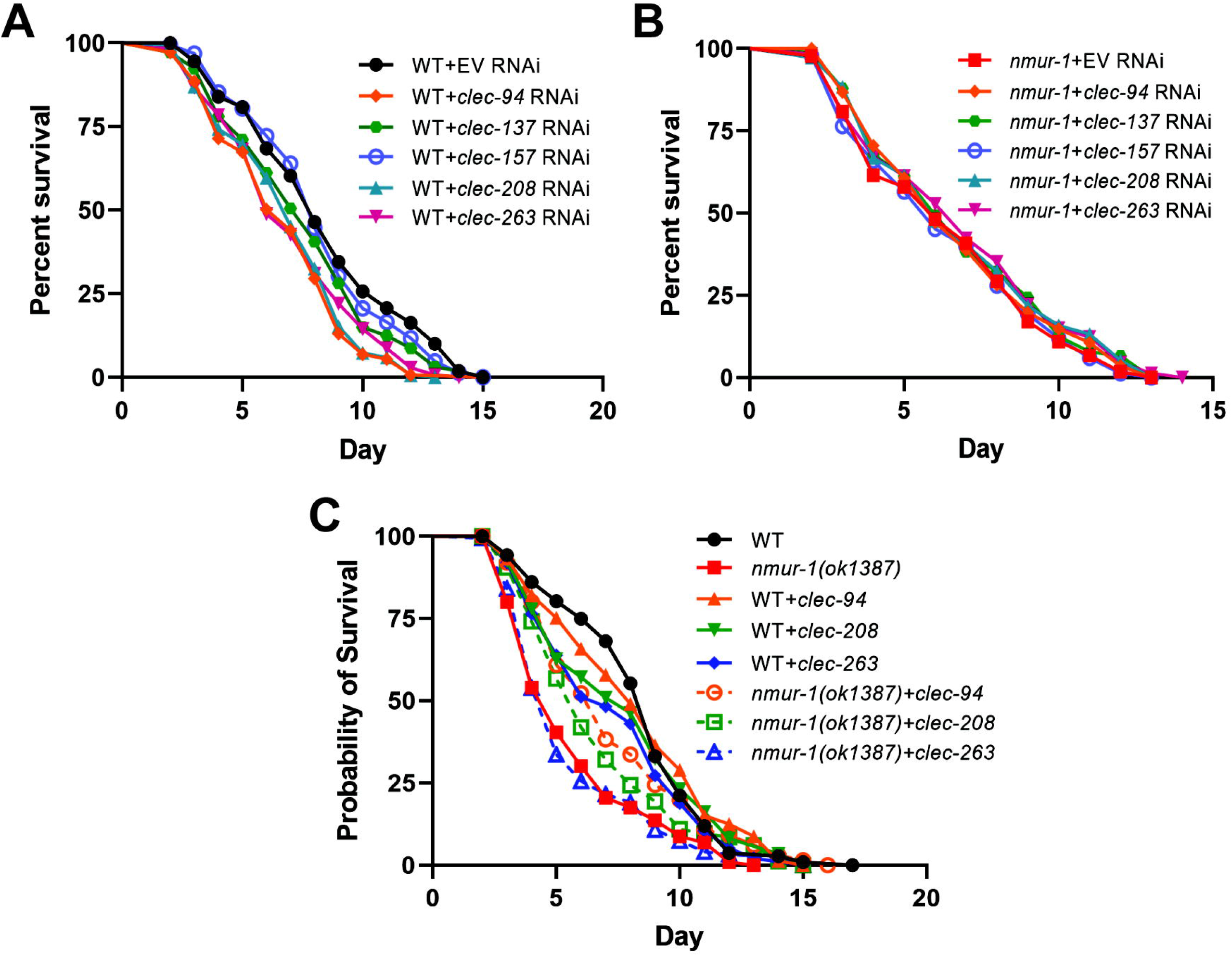
Functional loss of NMUR-1 differentially affects *C. elegans* survival against *E. coli* HT115, *E. faecalis* and *S. enterica*. Wild-type (WT) and *nmur-1(ok1387)* animals were exposed to *E. coli* HT115 **(A)**, *E. faecalis* **(B)**, or *S. enterica* **(C)** and scored for survival over time. Each graph is a combination of three independent experiments. Each experiment included *N* = 60 animals per strain. *P*-values represent the significance level of mutant survival relative to WT, *P* = 0.8706 in (A), *P* = 0.0004 in (B), and *P* < 0.0001 in (C).

There are many factors affecting the pathogenesis of the above-mentioned bacteria besides being Gram-negative or Gram-positive. For example, each pathogen can produce a unique set of virulence factors that promote pathogenicity through different mechanisms (Peterson, 1996). The diverse effects that NMUR-1 has on *C. elegans* survival suggest that it could have multiple physiological functions in the nematode’s defense against these bacteria. Given NMUR-1’s potential versatility in function, we sought to gain insights into NMUR-1’s functionality by focusing on its roles in defense against *E. faecalis* and *S. enterica*, two pathogens that have opposite effects on the survival of *nmur-1* mutant animals (reduced survival against *E. faecalis* but improved survival against *S. enterica* (Fig. 1). Below, we describe our study using these two pathogens as infecting agents.

### Pathogen avoidance behavior does not contribute to the altered survival of *nmur-1(ok1387)* animals exposed to *E. faecalis* or *S. enterica*

As described above, compared to wild-type animals, *nmur-1(ok1387)* animals exhibited altered survival following exposure to *E. faecalis* or *S. enterica* (Fig. 1). However, no differences in survival were observed between mutant and wild-type animals when they were fed worm food *E. coli* strain HT115 (Fig. 1), heat-killed *E. faecalis* (Fig. S2A), or heat-killed *S. enterica* (Fig. S2B). These results indicate that the *nmur-1* mutation affects *C. elegans* defense against living pathogens without impacting the nematode’s basic lifespan. The altered survival of mutant animals could be due to pathogen avoidance behavior, a known mechanism of *C. elegans* defense (Meisel and Kim, 2014). To test this possibility, we measured animal survival using full-lawn assays in which the agar plates were completely covered by pathogenic bacteria to eliminate pathogen avoidance. Consistent with our original (partial-lawn) survival assays, we observed that *nmur-1(ok1387)* animals exposed to *E. faecalis* died at a faster rate than wild-type controls, whereas mutants exposed to *S. enterica* died at a slower rate than wild-type controls (Fig. S3). These results suggest that lawn avoidance is not the reason for the altered survival observed in *nmur-1* mutants. We next conducted lawn occupancy behavioral assays to compare the magnitude of pathogen avoidance between wild-type and *nmur-1(ok1387)* animals. In these assays, a small lawn of pathogenic bacteria was seeded in the center of an agar plate, and a set number of synchronized young adult animals were placed on the lawn. The numbers of animals that stayed on and off the lawn were then counted at five time points over a period of 36 hours. Compared to wild-type animals, *nmur-1(ok1387)* animals overall did not show significant differences in the magnitudes of pathogen avoidance (Fig. 2). Taken together, these data suggest that pathogen avoidance behavior does not play a role in the altered survival of *nmur-1(ok1387)* animals exposed to *E. faecalis* or *S. enterica*.

**Fig. 2.**
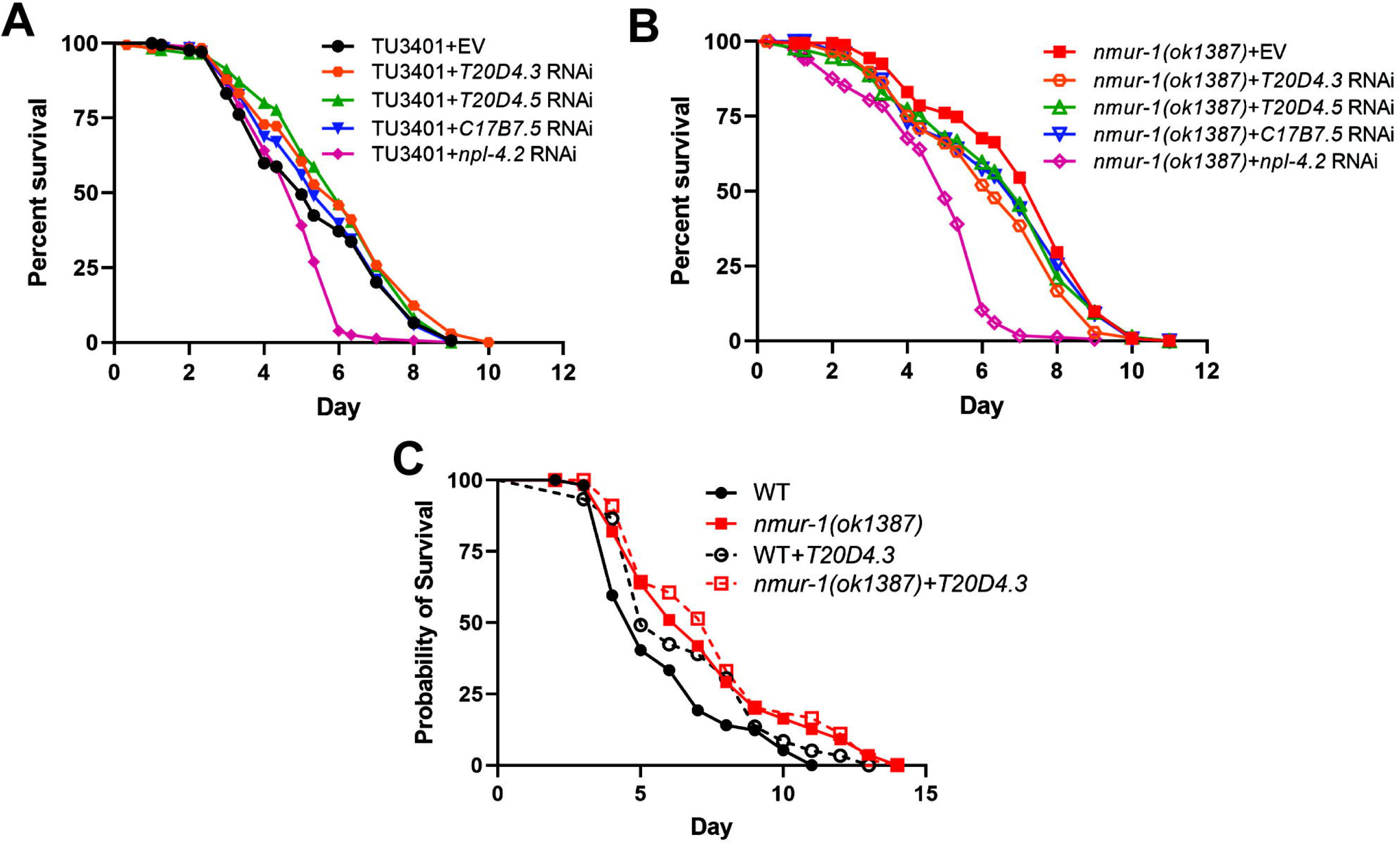
Functional loss of NMUR-1 does not affect pathogen avoidance behavior. WT and *nmur-1(ok1387)* animals were subjected to lawn occupancy assays in which the animals were placed on a small lawn of *E. faecalis* **(A)** or *S. enterica* **(B)** in a 3.5cm plate and monitored over time for their presence on the lawn. Each graph is a combination of three independent experiments. Each experiment included *N* = 60 animals per strain. Error bars represent the standard error of the mean (SEM).

### Functional loss of NMUR-1 changes intestinal accumulation of *E. faecalis* and *S. enterica*

Certain mutations in *C. elegans* can change pathogen accumulation in the intestine, leading to altered resistance to infection (Fuhrman et al., 2009). To examine whether the *nmur-1* mutation affects bacterial accumulation, we used GFP-tagged *E. faecalis* to visualize accumulated bacteria in the nematode’s intestine. Our results showed that the GFP fluorescence intensity in *nmur-1(ok1387)* animals was stronger than that seen in wild-type animals (Fig. 3A). Intestinal bacterial loads were quantified by counting the Colony Forming Units (CFU) of live bacterial cells recovered from the intestine. The mutant animals had more *E. faecalis* CFUs than wild-type animals (Fig. 3B). These results indicated that the *nmur-*1 mutation caused increased intestinal accumulation of *E. faecalis*. Such higher bacterial accumulation could be due to altered pathogen intake, expulsion, or clearance by immunity. These possible causes were further investigated. *C. elegans* feeds on bacterial food via rhythmic contractions (pumping) of its pharynx. To determine whether there were any differences in pathogen intake between wild-type and *nmur-1(ok1387)* animals, we measured their pharyngeal pumping rates on bacterial lawns. When feeding on *E. faecalis*, both wild-type and mutant animals showed similar pumping rates, reflecting similar levels of pathogen intake (Fig. 3C). Bacterial evacuation was assessed by measuring the defecation rate, defined as the time interval between expulsions of gut contents. We found that the defecation rate of *nmur-1(ok1387)* animals on *E. faecalis* was comparable to the rate observed in wild-type animals (Fig. 3D). Overall, these results indicate that the reduced survival of *nmur-1(ok1387)* animals exposed to *E. faecalis* is likely due to increased pathogen accumulation in the intestine. Because the *nmur-1* mutation did not alter *E. faecalis* intake or expulsion, the increased pathogen accumulation likely resulted from the mutants’ weakened innate immunity that was unable to clear the infection from the intestine.

**Fig 3.**
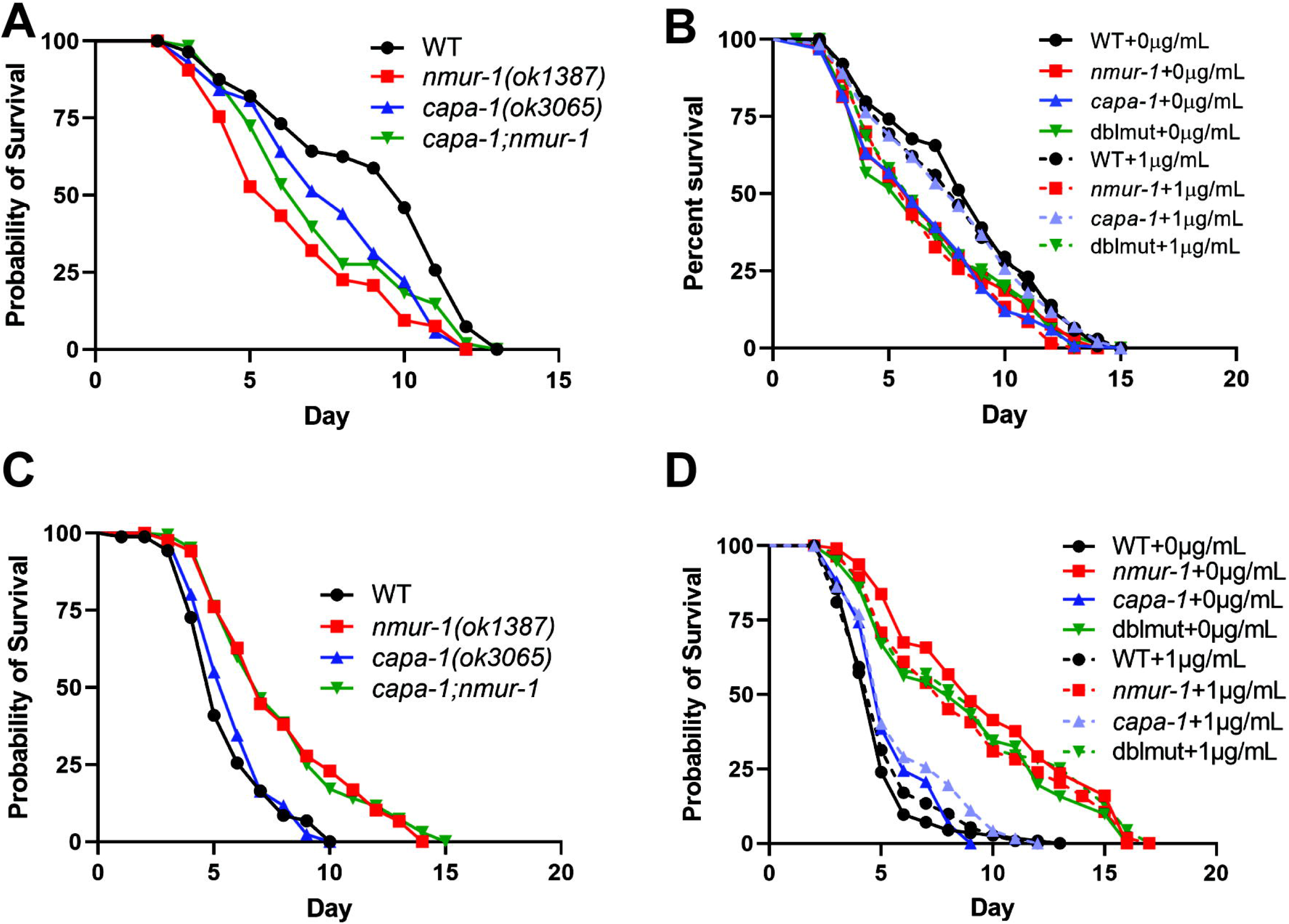
Functional loss of NMUR-1 changes intestinal accumulation of *E. faecalis* and *S. enterica*. **(A)** WT and *nmur-1(ok1387)* animals were exposed to GFP-expressing *E. faecalis* for 24 hours and then visualized using a Zeiss Axio Imager M2 fluorescence microscope. The scale bar indicates 500 µm. **(B)** WT and *nmur-1(ok1387)* animals were exposed to *E. faecalis* for 24 hours, and the Colony Forming Units (CFU) of live bacterial cells recovered from their intestines were quantified. The graph represents the combined results of three independent experiments. *N* = 10 animals per strain were used for each experiment. Error bars represent SEM. *P*-value represents the significance level of mutant gut colonization relative to WT, *P* = 0.0296. **(C)** WT and *nmur-1(ok1387)* animals were exposed to *E. faecalis* for 24 hours. Pharyngeal pumping rates of animals were counted as the number of pumps per 30 sec. Counting was conducted in triplicate and averaged to give a pumping rate. The measurements included *N* = 10 adult animals per strain. Error bars represent SEM. *P* = 0.1068. **(D)** WT and *nmur-1(ok1387)* animals were exposed to *E. faecalis* for 24 hours. Defecation rates of animals were measured as the average time of 10 intervals between two defecation cycles. The measurements included *N* = 10 animals per strain. *P* = 0.1379. **(E)** WT and *nmur-1(ok1387)* animals were exposed to GFP-expressing *S. enterica* for 24 hours and then visualized using a Zeiss Axio Imager M2 fluorescence microscope. The scale bar indicates 500 µm. **(F)** WT and *nmur-1(ok1387)* animals were exposed to GFP-expressing *S. enterica* for 24 hours, and the CFU of live bacterial cells recovered from their intestines were quantified. The graph represents the combined results of three independent experiments. *N* = 10 animals per strain were used for each experiment. Error bars represent SEM. *P* = 0.0290. **(G)** WT and *nmur-1(ok1387)* animals were exposed to *S. enterica* for 24 hours. Pharyngeal pumping rates of animals were counted as the number of pumps per 30 sec. Counting was conducted in triplicate and averaged to give a pumping rate. The measurements included *N* = 10 adult animals per strain. Error bars represent SEM. *P* = 0.0006. **(H)** WT and *nmur-1(ok1387)* animals were exposed to *S. enterica* for 24 hours. Defecation rates of animals were measured as the average time of 10 intervals between two defecation cycles. The measurements included *N* = 10 animals per strain. *P* = 0.6310.

We next performed similar experiments to investigate intestinal accumulation, intake, and expulsion of *S. enterica*. Our results showed that, compared to wild-type animals, *nmur-1(ok1387)* animals exhibited slightly weaker GFP-tagged *S. enterica* fluorescence intensities (Fig. 3E) and had significantly lower CFU counts (Fig. 3F), indicating that the *nmur-*1 mutation decreased intestinal accumulation of *S. enterica*. Mutant animals also had slightly higher pharyngeal pumping rates than wild-type animals (Fig. 3G), suggesting that pathogen intake was increased in the mutants despite their reduced bacterial accumulation. The defecation rate of *nmur-1(ok1387)* animals on *S. enterica* was comparable to that of wild-type animals (Fig. 3H). These results indicate that the enhanced survival of *nmur-1(ok1387)* animals exposed to *S. enterica* is likely due to decreased pathogen accumulation in the intestine. This decrease did not result from reduced pathogen intake (pathogen intake actually improved) or increased expulsion but likely resulted from enhanced innate immunity in the mutants that cleared the infection from the intestine. Taken together, both the *E. faecalis* and *S. enterica* studies suggest that NMUR-1 could regulate innate immunity in *C. elegans* defense against pathogen infection but plays opposite roles in such potential regulation against the two pathogens.

### NMUR-1 regulates *C. elegans* transcriptional activity

To gain insights into how NMUR-1 regulates *C. elegans* defense at the molecular level, we employed RNA sequencing to profile gene expression in wild-type and *nmur-1(ok1387)* animals with or without pathogen infection. For these studies, we collected RNA samples from wild-type and *nmur-1(ok1387)* animals with or without exposure to *E. faecalis* or *S. enterica* (five replicates per group). The samples were then submitted to the Washington State University Genomics Core for RNA-seq analysis. The resulting sequence data (FASTQ files) were deposited in the National Center for Biotechnology Information (NCBI) Sequence Read Archive (SRA) database through the Gene Expression Omnibus (GEO). The processed gene quantification files and differential expression files were deposited in the GEO. All of these data can be accessed through the GEO with the accession number GSE154324.

Using the resulting sequencing data, we first compared the gene expression profiles of *nmur-1(ok1387)* animals with those of wild-type animals under normal conditions (i.e., without infection). In total, 14,206 genes were detected and quantified with a false discovery rate (FDR) of 5%. Among these genes, 1,599 were upregulated and 275 were downregulated at least two-fold in *nmur-1(ok1387)* animals relative to wild-type animals. The change in expression of such a large number of genes in the mutant worms indicates that the lack of NMUR-1 profoundly affects gene expression in *C. elegans*. Gene ontology (GO) analysis of the 1,599 upregulated genes identified 26 significantly enriched molecular functions, including protein kinase/phosphatase activity, ribonucleotide binding, and cuticle structure activity (Table S1). Interestingly, GO analysis of the 276 downregulated genes showed 11 significantly enriched molecular functions, all of which involve transcription-related DNA binding activities (Table 1A). Twenty-three downregulated genes possess these activities and contribute to the functional enrichment, most of which encode transcription factors involved in binding to RNA polymerase II regulatory regions (Table 1B). These results indicate that NMUR-1 could affect transcription by regulating the expression of these transcription factors.

**Table 1.**
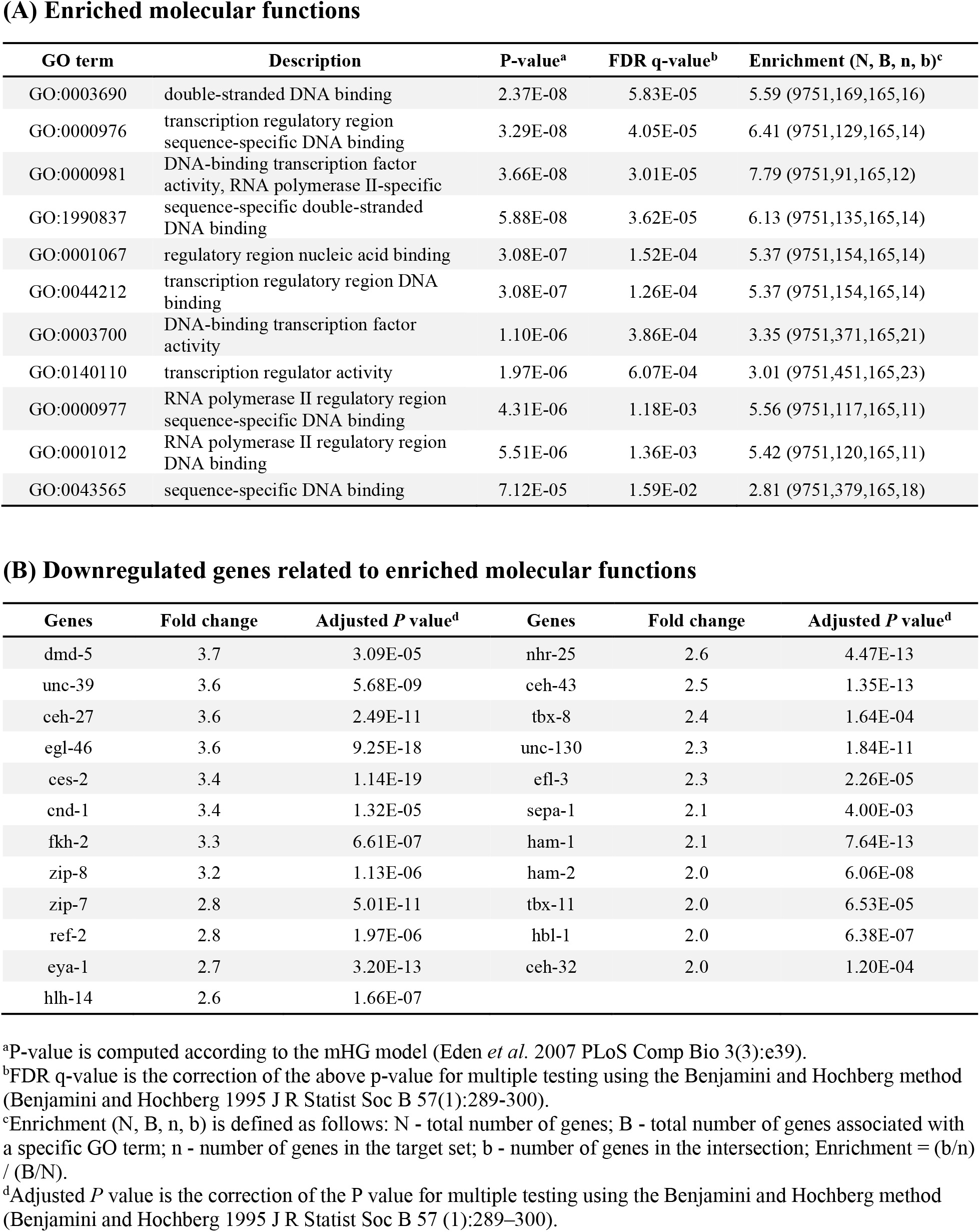
Gene ontology (GO) analysis of downregulated genes in *nmur-1(ok1387)* animals relative to wild-type animals.

Because of the high importance of transcription in the fundamental functions of life and the potential regulatory role of NMUR-1 in this process, we next examined how NMUR-1, or the lack thereof, impacts transcription. To this end, we developed a novel *in vitro* transcription system to measure *C. elegans* transcriptional activity (Wibisono et al., 2020). One of the biggest obstacles for biochemical studies of *C. elegans* is the high difficulty of preparing functionally active nuclear extract due to its thick surrounding cuticle. In our transcription system, Balch homogenization was employed to effectively disrupt worms, and functionally active nuclear extracts were obtained through subcellular fractionation. Subsequently, the nuclear extracts were used to re-constitute *in vitro* transcription reactions with a linear DNA template containing the worm *Δpes-10* promoter, followed by PCR-based, non-radioactive detection of transcripts (Wibisono *et al*., 2020). Using this system, we were able to compare the general transcriptional activity of *nmur-1(ok1387)* animals with that of wild-type animals at the whole-organism level. PCR and gel analysis of the transcript products showed substantial increases in transcriptional activity in reactions with mutant nuclear extract compared to those with wild-type nuclear extract (Fig. 4A). To quantitatively compare the transcriptional activity of wild-type and mutant nuclear extracts, we next titrated the DNA template with varying amounts of each extract and used qRT-PCR to quantify transcripts at each titration point. The resulting data were then fitted with the Michaelis-Menten model and the non-linear least-squares method, as we described previously (Wibisono *et al*., 2020). The titration curves and two titration parameters, namely, the maximum yield and the amount of nuclear extract needed to reach 50% of the maximum yield, are shown in Fig. 4B. Compared to the titrations with wild-type nuclear extract, those with mutant extract yielded 2.33-fold maximum number of transcripts and only needed 0.67-fold extract to reach 50% of the maximum yield (Fig. 4B), indicating that the reactions performed with mutant extract were more robust and had higher transcriptional activity. These results were surprising given the fact that the expression of many transcription factors was downregulated in the mutant animals. Taken together, our data suggest that NMUR-1 suppresses transcriptional activity in wild-type animals and that functional loss of NMUR-1 abrogates this suppression, which explains why a large number of genes (1,599 genes) were upregulated in *nmur-1(ok1387)* animals relative to wild-type animals.

**Fig. 4.**
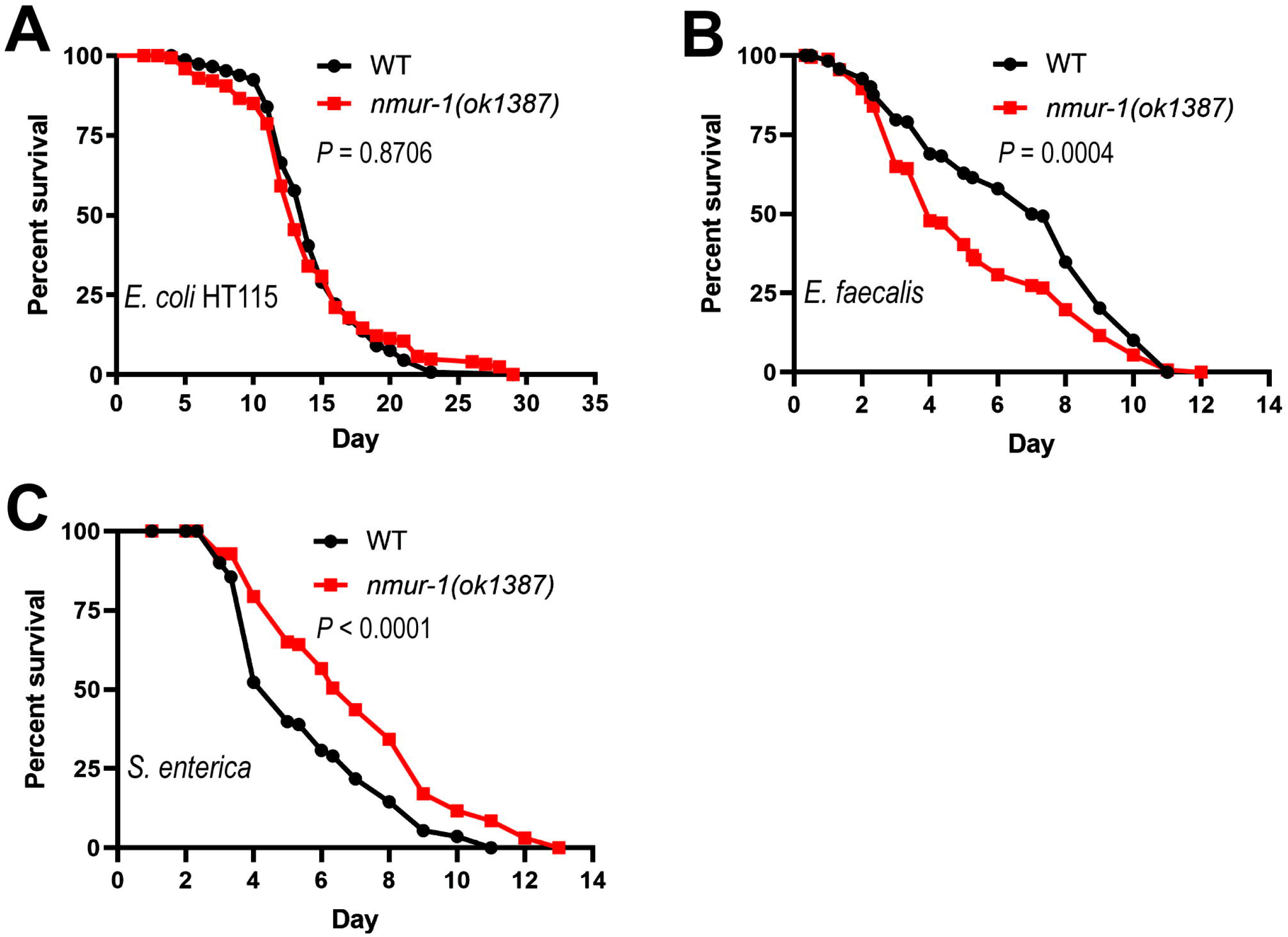
NMUR-1 regulates transcriptional activity. **(A)** *In vitro* transcription assays were performed using nuclear extract from WT or *nmur-1(ok1387)* animals. After transcription, RNA was purified, reverse transcribed, and amplified by PCR, followed by gel analysis. Lane 1, transcription with WT nuclear extract; lane 2, transcription with *nmur-1(ok1387)* nuclear extract; lane 3, transcriptional positive control (with HeLa nuclear extract); lane 4, transcriptional negative control (without any nuclear extract); lane 5, PCR positive control; lane 6, PCR negative control (without DNA template). **(B)** WT and mutant nuclear extracts were titrated using the *in vitro* transcription system. The RNA copy number generated from transcription was plotted against the amount of nuclear extract used in the reaction, followed by fitting with the Michaelis-Menten model and the non-linear least-squares method to generate the titration curves and calculate titration parameters. **(C)** qRT-PCR analysis of GFP expression after two-hour heat shock in *CL2070(dvls70)* and *nmur-1(ok1387);CL2070(dvls70)* animals. Asterisk (*) denotes a significant difference between *CL2070(dvls70)* and *nmur-1(ok1387);CL2070(dvls70)* animals.

We also employed *in vivo* transcription assays to evaluate how the lack of NMUR-1 affects transcriptional activity in *C. elegans*. For these assays, transgenic strain CL2070, which contains the *hsp-16.2p::GFP* reporter transgene, was used. This strain does not express detectable GFP under standard conditions but broadly expresses GFP in all tissues after heat shock, allowing for the assessment of overall transcriptional activity in living animals (Link et al., 1999). The CL2070 animals were outcrossed with our wild-type animals and then crossed with *nmur-1(ok1387)* animals. The resulting animals (*CL2070(dvls70)* and *CL2070(dvls70);nmur-1(ok1387))* were grown to the L4 larval stage, heat shocked at 35°C for 2 hours, and then returned to 20°C for GFP mRNA quantification by qRT-PCR. Our results showed that GFP expression was significantly increased in *CL2070(dvls70);nmur-1(ok1387)* animals compared to *CL2070(dvls70)* animals (Fig. 4C), indicating that the *nmur-1* mutation causes higher overall transcriptional activity, which is consistent with the results of our *in vitro* transcription assays.

### NMUR-1 regulates the innate immune response to *E. faecalis* by controlling the expression of C-type lectins

Next, we asked how NMUR-1 regulates gene expression in *C. elegans* in response to *E. faecalis* infection. By examining the gene expression profiles of *nmur-1(ok1387)* animals relative to wild-type animals exposed to *E. faecalis*, we found that 229 genes were upregulated and 106 genes were downregulated at least two-fold. GO analysis of the upregulated genes identified 11 significantly enriched biological processes, all of which involve defense responses to bacteria or other external stimuli (Table S2). This is consistent with the fact that *E. faecalis* is a pathogenic agent for *C. elegans* (Yuen and Ausubel, 2018). GO analysis of the downregulated genes yielded one significantly enriched molecular function involving carbohydrate binding activity (Table 2A). Fifteen of the downregulated genes were related to this activity, 14 of which encode C-type lectins (Table 2B). C-type lectins are a group of proteins that contain one or more C-type lectin-like domains (CTLDs) and recognize complex carbohydrates on cells and tissues (Drickamer and Dodd, 1999; Takeuchi et al., 2008). C-type lectins may act in pathogen recognition or function as immune effectors to fight infection (Pees *et al*., 2016; Sun *et al*., 2016). Therefore, downregulation of C-type lectins in *nmur-1(ok1387)* animals might explain the reduced survival observed in mutant animals following exposure to *E. faecalis*.

**Table 2.** GO analysis of downregulated genes in *nmur-1(ok1387)* animals relative to wild-type animals exposed to *E. faecalis*.

**(A) Enriched molecular functions**
| GO term | Description | P-value<sup>a</sup> | FDR q-value<sup>b</sup> | Enrichment (N, B, n, b)<sup>c</sup> |
| --- | --- | --- | --- | --- |
| GO:0030246 | carbohydrate binding | 2.29E-15 | 5.65E-12 | 17.45 (10161,182,48,15) |

**(B) Downregulated genes related to the enriched molecular functions**
| Genes | Fold change | Adjusted *P* value<sup>d</sup> | Genes | Fold change | Adjusted *P* value<sup>d</sup> |
| --- | --- | --- | --- | --- | --- |
| clec-207 | 5.4 | 4.16E-15 | clec-161 | 2.4 | 1.10E-04 |
| clec-94 | 4.9 | 1.50E-12 | clec-95 | 2.4 | 2.91E-04 |
| clec-137 | 4.3 | 3.68E-10 | clec-99 | 2.4 | 6.31E-05 |
| clec-263 | 3.7 | 1.68E-08 | clec-197 | 2.4 | 7.91E-04 |
| clec-208 | 3.4 | 2.01E-07 | clec-136 | 1.9 | 2.03E-02 |
| clec-157 | 2.9 | 3.25E-06 | clec-174 | 1.9 | 2.50E-03 |
| pqn-84 | 2.8 | 2.76E-06 | clec-190 | 1.8 | 1.29E-04 |
| clec-219 | 2.5 | 2.92E-04 |  |  |  |

<sup>a</sup>P-value is computed according to the mHG model (Eden *et al.* 2007 PLoS Comp Bio 3(3):e39).
<sup>b</sup>FDR q-value is the correction of the above p-value for multiple testing using the Benjamini and Hochberg method (Benjamini and Hochberg 1995 J R Statist Soc B 57(1):289-300).
<sup>c</sup>Enrichment (N, B, n, b) is defined as follows: N - total number of genes; B - total number of genes associated with a specific GO term; n - number of genes in the target set; b - number of genes in the intersection; Enrichment = (b/n) / (B/N).
<sup>d</sup>Adjusted *P* value is the correction of the *P* value for multiple testing using the Benjamini and Hochberg method (Benjamini and Hochberg 1995 J R Statist Soc B 57 (1):289–300).

To determine whether NMUR-1-regulated C-type lectin genes contribute to the reduced survival of *nmur-1(ok1387)* animals, we inactivated these genes using RNAi-mediated knockdown in wild-type and *nmur-1(ok1387)* animals and assayed for their survival against *E. faecalis*-mediated killing. Five of the most downregulated C-type lectin genes (*clec-94, clec-137, clec-157, clec-208,* and *clec-263*) were individually targeted in these studies. While RNAi of these genes did not further reduce the survival of *nmur-1(ok1387)* animals (Fig. 5B), knockdown of four of the five genes (all except *clec-157*) significantly suppressed the survival of wild-type animals (Fig. 5A). These results indicate that some of the NMUR-1-regulated C-type lectin genes are indeed required for *C. elegans* defense against *E. faecalis* infection.

**Fig. 5.**
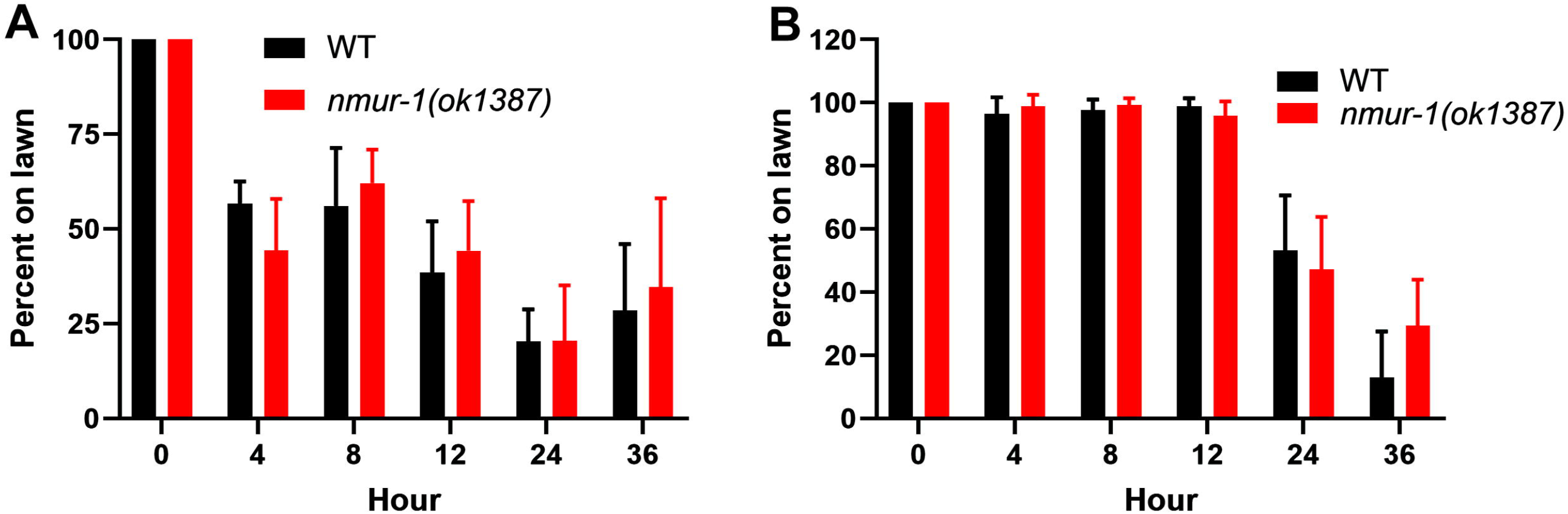
NMUR-1-regulated C-type lectin genes are required for *C. elegans* defense against *E. faecalis*. **(A)** WT and **(B)** *nmur-1(ok1387)* animals grown on dsRNA for C-type lectin genes or the empty vector (EV) control were exposed to *E. faecalis* and scored for survival over time. The graphs are a combination of three independent replicates. Each experiment included *N* = 60 animals per strain. *P*-values in (A) represent the significance level of RNAi treatment relative to WT+EV: WT+*clec-94*, *P* < 0.0001; WT+*clec-137*, *P* = 0.0220; WT+*clec-157*, *P* = 0.3728; WT+*clec-208*, *P* < 0.0001; WT+*clec-263*, *P* < 0.0001. *P*-values in (B) represent the significance level of the RNAi treatment relative to *nmur-1(ok1387)*+EV: *nmur-1(ok1387)*+*clec-94*, *P* = 0.3633; *nmur-1(ok1387)*+*clec-137*, *P* = 0.1841; *nmur-1(ok1387)*+*clec-157*, *P* = 0.8617; *nmur-1(ok1387)*+*clec-208*, *P* = 0.1348; *nmur-1(ok1387)*+*clec-263*, *P* = 0.1320. **(C)** WT and *nmur-1(ok1387)* animals in which C-type lectin expression was rescued were exposed to *E. faecalis* and scored for survival over time. All C-type lectin genes were rescued using their own promoters. The graph is a combination of three independent experiments. Each experiment included *N* = 60 animals per strain. *P*-values represent the significance of the survival of rescue strains relative to *nmur-1(ok1378)* animals: WT, *P* < 0.0001; *nmur-1(ok1387)*+*clec-94*, *P* < 0.0001; *nmur-1(ok1387)*+*clec-208*, *P* = 0.0025; *nmur-1(ok1387)*+*clec-263*, *P* = 0.9799.

Since knockdown of C-type lectins in wild-type animals recaptured the reduced survival phenotype of *nmur-1(ok1387)* animals, we next asked whether complementing the expression of these C-type lectins in mutant animals could rescue their survival phenotype. To this end, we employed transgenes to express *clec-94*, *clec-208*, and *clec-263*, respectively, in *nmur-1(ok1387)* and wild-type animals. Survival assays against *E. faecalis* showed that while overexpression of these C-type lectins had no effect on wild-type animal survival, the expression of *clec-94* or *clec-208* partially rescued the reduced survival of the mutant animals (Fig. 5C), confirming that the downregulation of C-type lectins contributes to the reduced survival of *nmur-1* mutants challenged with *E. faecalis*.

### NMUR-1 suppresses the innate immune response to *S. enterica* by inhibiting unfolded protein response (UPR) genes

We also compared the gene expression profiles of *nmur-1(ok1387)* animals with those of wild-type animals exposed to *S. enterica* and found that 28 genes were upregulated and 28 genes were downregulated at least two-fold. While GO analysis of the downregulated genes did not yield any enriched GO terms, a similar analysis of the upregulated genes identified four significantly enriched biological processes and two significantly enriched molecular functions (Table 3A). Among these enrichments, the biological process “protein quality control for misfolded or incompletely synthesized proteins” was the most significantly enriched (FDR = 7.89E-5) and has a known role in immune responses to pathogen infection (Grootjans et al., 2016). Under an environmental stress, such as pathogen infection, increased demand for protein folding causes ER stress, which activates the UPR pathways to alleviate the stress and maintain protein homeostasis. This process is known to be controlled by the nervous system (Aballay, 2013). Four of the upregulated genes (*T20D4.3*, *T20D4.5*, *C17B7.5*, and *npl-4.2*) are related to UPR activity and contributed to the biological process enrichment (Table 3B).

**Table 3.** Enrichment of biological processes and molecular functions revealed by GO analysis of upregulated genes in *nmur-1(ok1387)* animals relative to wild-type animals exposed to *S. enterica*.

**(A) Enriched biological processes and molecular functions**
| GO term | Description | P-value<sup>a</sup> | FDR q-value<sup>b</sup> | Enrichment (N, B, n, b)<sup>c</sup> |
| --- | --- | --- | --- | --- |
| **<u>Biological process</u>** |  |  |  |  |
| GO:0006515 | protein quality control for misfolded or incompletely synthesized proteins | 1.35E-8 | 7.89E-5 | 137.71 (9812,19,15,4) |
| GO:0006516 | glycoprotein catabolic process | 3.45E-7 | 1.01E-3 | 196.24 (9812,10,15,3) |
| GO:0006517 | protein deglycosylation | 1.04E-6 | 2.03E-3 | 140.17 (9812,14,15,3) |
| GO:0009100 | glycoprotein metabolic process | 2E-5 | 2.92E-2 | 54.51 (9812,36,15,3) |
| **<u>Molecular function</u>** |  |  |  |  |
| GO:0000224 | peptide-N4-(N-acetyl-beta-glucosaminyl)asparagine amidase activity | 2.41E-7 | 5.96E-4 | 218.04 (9812,9,15,3) |
| GO:0016811 | hydrolase activity, acting on carbon-nitrogen (but not peptide) bonds, in linear amides | 2.98E-5 | 3.67E-2 | 47.86 (9812,41,15,3) |

**(B) Upregulated genes related to the enriched biological process protein quality control**
| Genes | Fold change | Adjusted *P* value<sup>d</sup> |
| --- | --- | --- |
| C17B7.5 | 2.9 | 2.57E-14 |
| T20D4.5 | 2.8 | 2.40E-15 |
| npl-4.2 | 2.8 | 1.22E-22 |
| T20D4.3 | 2.6 | 1.60E-12 |

<sup>a</sup>P-value is computed according to the mHG model (Eden *et al.* 2007 PLoS Comp Bio 3(3):e39).
<sup>b</sup>FDR q-value is the correction of the above p-value for multiple testing using the Benjamini and Hochberg method (Benjamini and Hochberg 1995 J R Statist Soc B 57(1):289-300).
<sup>c</sup>Enrichment (N, B, n, b) is defined as follows: N - total number of genes; B - total number of genes associated with a specific GO term; n - number of genes in the target set; b - number of genes in the intersection; Enrichment = (b/n) / (B/N).
<sup>d</sup>Adjusted *P* value is the correction of the *P* value for multiple testing using the Benjamini and Hochberg method (Benjamini and Hochberg 1995 J R Statist Soc B 57 (1):289–300).

To determine whether these NMUR-1-regulated UPR genes play any roles in the enhanced survival of *nmur-1(ok1387)* animals, we inactivated these genes individually using RNAi and then assayed the animals for survival against *S. enterica*. Because these genes are mainly expressed in neurons (WormBase), we employed strain *TU3401*, which is capable of RNAi-mediated gene knockdown in neurons (Calixto et al., 2010), in our experiments. While RNAi of *npl-4.2* suppressed the survival of both TU3401 and *nmur-1(ok1387);TU3401* animals following exposure to *S. enterica*, RNAi silencing of *T20D4.3* specifically suppressed the enhanced survival of *nmur-1(ok1387);TU3401* animals (Fig. 6A and 6B). In comparison, RNAi of *T20D4.5* or *C17B7.5* had no effect on animal survival (Fig. 6A and 6B). The two genes, *npl-4.2* and *T20D4.3*, that showed effects on survival have known roles in the UPR. *npl-4.2* has been implicated in ER-associated protein degradation. Specifically, NPL-4 forms a complex with CDC-48 ATPase and UFD-1, which extracts polyubiquitin-tagged misfolded proteins from the ER membrane and facilitate their degradation by the proteasome (Bays et al., 2001; Ye et al., 2001). *T20D4.3* encodes a peptide N-glycanase and acts in ER-associated degradation (ERAD) system to surveil ER quality control/homeostasis for newly synthesized glycoproteins; it reportedly functions in longevity, dauer formation, and pathogen responses (Gaglia et al., 2012).

**Fig. 6.**
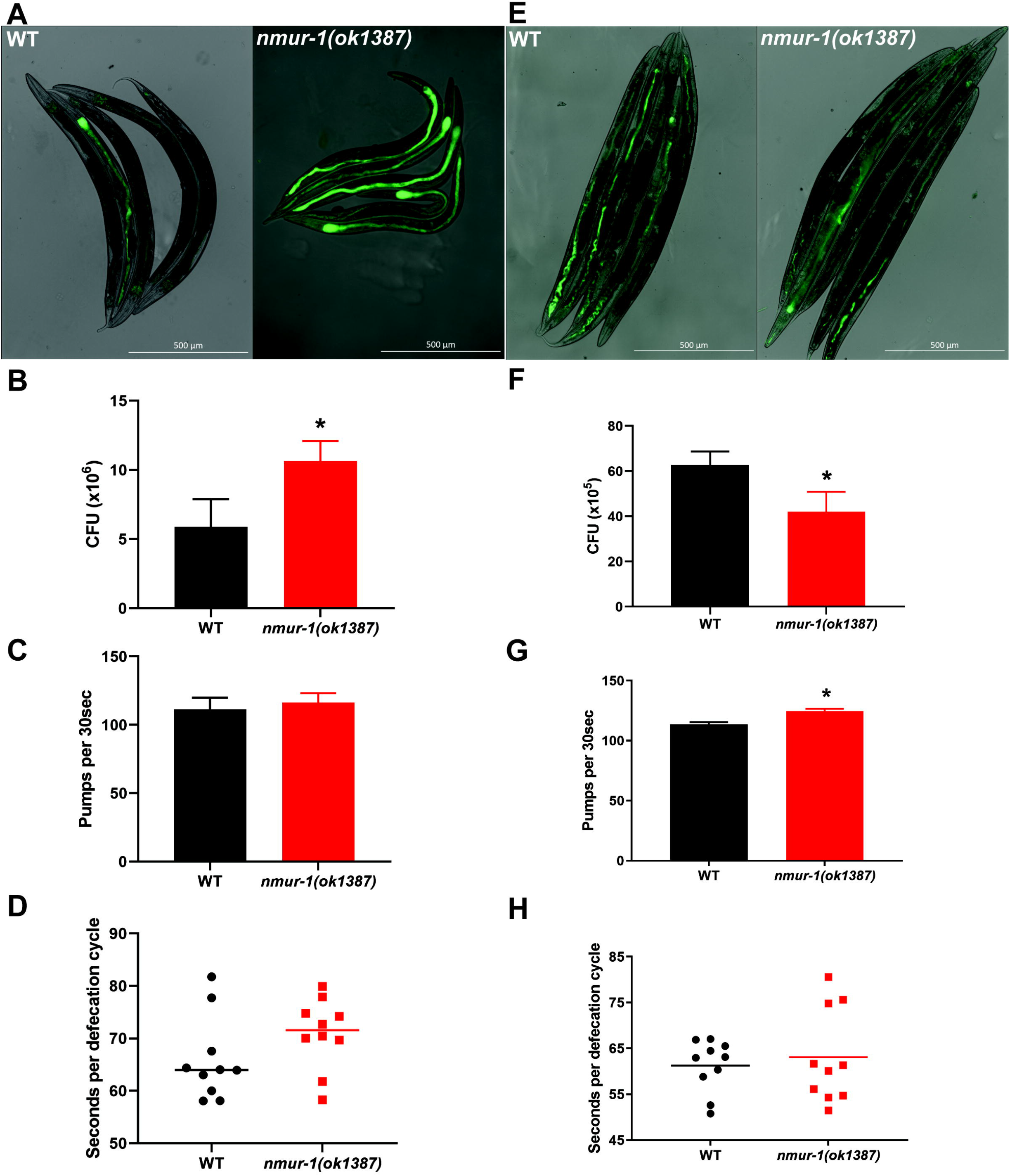
NMUR-1-regulated UPR genes are required for *C. elegans* defense against *S. enterica*. **(A)** TU3401(*uIs69*) and (**B)** *nmur-1(ok1387);*TU3401 animals grown on dsRNA for UPR genes or the empty vector (EV) control were exposed to *S. enterica* and scored for survival over time. The graphs are a combination of three independent replicates. Each experiment included *N* = 60 animals per strain. *P-*values in (A) represent the significance level of the RNAi treatment relative to the TU3401+EV animals: TU3401+*T20D4.3, P* = 0.4224; TU3401+*T20D4.5, P* < 0.6102; TU3401+*C17B7.5, P* = 0.5905; TU3401+*npl-4.2, P* < 0.0001. *P*-values in (B) represent the significance level of RNAi treatment relative to the *nmur-1(ok1387);*TU3401+EV animals: *nmur-1(ok1387);*TU3401+*T20D4.3, P* = 0.0007; *nmur-1(ok1387);*TU3401+*T20D4.5, P*= 0.0993; *nmur-1(ok1387);*TU3401+*C17B7.5, P* = 0.1216; *nmur-1(ok1387);*TU3401+*npl-4.2, P* < 0.0001. **(C)** WT, *nmur-1(ok1387),* and *T20D4.3* overexpression animals were exposed to *S. enterica. T20D4.3* was overexpressed using its own promoter. The graph is a combination of three independent experiments. Each experiment included *N =* 60 animals per strain. *P-*values represent the significance overexpression survival relative to the WT animals: *nmur-1(ok1387)*, *P* = 0.0031; WT+*T20D4.3, P =* 0.0336*; nmur-1(ok1387)*+*T20D4.3, P <* 0.0001*. P-*values represent the significance overexpression survival relative to the *nmur-1(ok1387)* animals: WT+*T20D4.3, P =* 0.2826; *nmur-1(ok1387)+T20D4.3, P =* 0.5181.

Because up-regulation of the UPR genes in *nmur-1(ok1387)* animals leads to enhanced survival of the mutants, we asked whether overexpression of these genes in wild-type animals can mimic the enhanced survival phenotype. To this end, we generated transgenic strains JRS78 and JRS79 that overexpress *T20D4.3* mRNA in wild-type *N2* and *nmur-1(ok1387)* animals, respectively (Table S3). Survival assays against *S. enterica* showed that overexpression of *T20D4.3* significantly improved the survival of wild-type animals but did not further increase the survival of *nmur-1* mutant animals (Fig. 6C). Taken together, our results indicate that some NMUR-1-regulated UPR genes are important for *C. elegans* defense against *S. enterica*, and that NMUR-1 can suppress the innate immune response to *S. enterica* by inhibiting UPR genes.

### The NMUR-1 ligand CAPA-1 is required for *C. elegans* defense against *E. faecalis* but not against *S. enterica*

CAPA-1, a homolog of vertebrate NMU and insect *capability* peptides, is an endogenous ligand of NMUR-1 (Lindemans et al., 2009; Watteyne et al., 2020). To determine whether CAPA-1 plays any roles in *C. elegans* defense, we examined the survival of a loss-of-function mutant of CAPA-1, *capa-1(ok3065)*, after *E. faecalis* exposure. Compared to wild-type animals, *capa-1(ok3065)* animals showed reduced survival, a phenotype also displayed by *nmur-1(ok1387)* animals (Fig. 7A). We then crossed *capa-1(ok3065)* with *nmur-1(ok1387)* animals to generate *capa-1(ok3065);nmur-1(ok1387)* double mutants. These double mutants exhibited reduced survival against *E. faecalis*, similar to *nmur-1(ok1387)* animals (Fig. 7A). These results indicate that CAPA-1 and NMUR-1 likely function in the same pathway to mediate *C. elegans* defense against *E. faecalis*. To investigate whether CAPA-1 is a ligand of NMUR-1 in the defense response, we tested whether exogenous administration of a synthetic CAPA-1 peptide could rescue the reduced survival phenotype of the mutant animals. To this end, we exposed wild-type and mutant animals to *E. faecalis* on BHI media containing 1 µg/mL synthetic CAPA-1 peptide and scored their survival over time. Our results showed that exogenous CAPA-1 peptide rescued the reduced survival phenotype of *capa-1(ok3065)* animals but failed to rescue *nmur-1(ok1387)* animals or *capa-1(ok3065);nmur-1(ok1387)* double mutants (Fig. 7B). These results demonstrate that the function of NMUR-1 in defense depends on the presence of CAPA-1, and *vice versa*, indicating that CAPA-1/NMUR-1 function as a ligand/receptor pair in mediating *C. elegans* defense against *E. faecalis*.

**Fig. 7.**
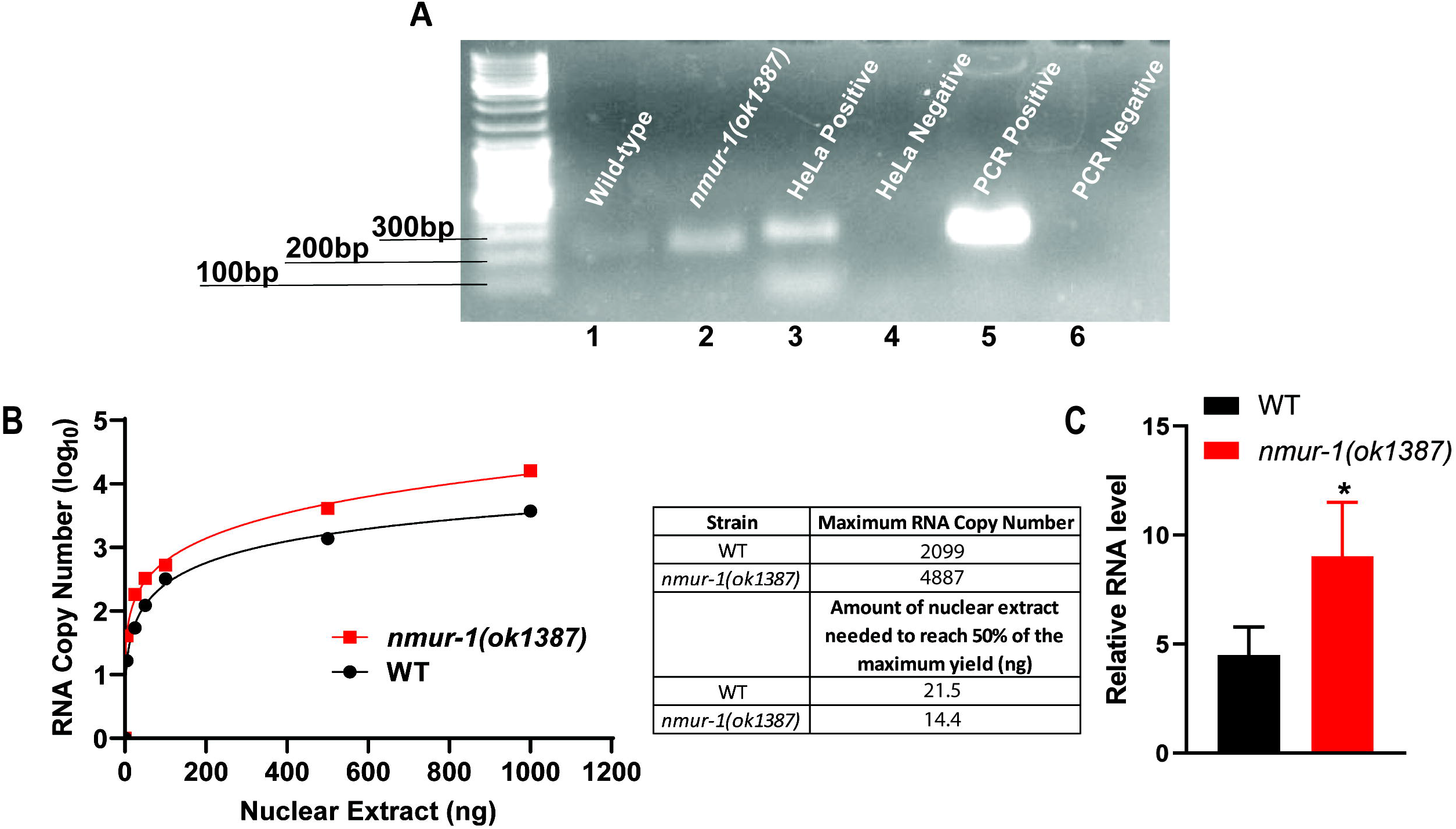
CAPA-1 is required for *C. elegans* defense against *E. faecalis* but not *S. enterica*. **(A)** WT, *nmur-1(ok1387), capa-1(ok3065),* and *capa-1(ok3065);nmur-1(ok1387)* animals were exposed to *E. faecalis* and scored for survival over time. The graph is a combination of three independent experiments. Each experiment included *N =* 60 animals per strain. *P*-values represent the significance level of the mutant survival relative to the WT: *nmur-1(ok1387), P* = 0.0005); *capa-1(ok3065), P* = 0.0134; *capa-1(ok3065);nmur-1(ok1387), P* = 0.0028. **(B)** WT, *nmur-1(ok1387), capa-1(ok3065),* and dblmut (*capa-1(ok3065);nmur-1(ok1387)*) animals were exposed to *E. faecalis* on plates contain either 0 or 1µg of synthetic CAPA-1 peptide and scored for survival over time. The graph is a combination of three independent experiments. Each experiment included *N =* 60 animals per strain. *P*-values represent the significance level of the survival for WT+0µg/mL vs. WT+1µg/mL: *P = 0.7467*; *nmur-1(ok1387)*+0µg/mL vs. *nmur-1(ok1387)*+1µg/mL: *P = 0.2515*; *capa-1(ok3065)+*0µg/mL vs. *capa-1(ok3065)+*1µg/mL: *P = 0.0005*; *capa-1(ok3065);nmur-1(ok1387)+*0µg/mL vs. *capa-1(ok3065);nmur-1(ok1387)+*1µg/mL: *P = 0.8313.* **(C)** WT, *nmur-1(ok1387), capa-1(ok3065),* and *capa-1(ok3065);nmur-1(ok1387)* animals were exposed to *S. enterica* and scored for survival over time. The graph is a combination of three independent experiments. Each experiment included *N =* 60 animals per strain. *P*-value represents the significance level of the mutant survival relative to the WT: *nmur-1(ok1387, P* < 0.0001; *capa-1(ok3065), P* = 0.2323; *capa-1(ok3065);nmur-1(ok1387), P <* 0.0001. **(D)** WT, *nmur-1(ok1387), capa-1(ok3065),* and dblmut (*capa-1(ok3065);nmur-1(ok1387)*) animals were exposed to *S. enterica* on plates contain either 0 or 1µg of synthetic CAPA-1 peptide and scored for survival over time. The graph is a combination of three independent experiments. Each experiment included *N =* 60 animals per strain. *P*-values represent the significance level of the survival for WT+0µg/mL vs. WT+1µg/mL: *P = 0.6683*; *nmur-1(ok1387)*+0µg/mL vs. *nmur-1(ok1387)*+1µg/mL: *P = 0.1701*; *capa-1(ok3065)+*0µg/mL vs. *capa-1(ok3065)+*1µg/mL: *P = 0.0779*; *capa-1(ok3065);nmur-1(ok1387)+*0µg/mL vs. *capa-1(ok3065);nmur-1(ok1387)+*1µg/mL: *P = 0.3087.*

To determine whether CAPA-1 also functions in the *C. elegans* defense response to *S. enterica*, we subjected wild-type and mutant animals to survival assays against *S. enterica*. Results showed that while *capa-1(ok3065)* animals displayed survival at the wild-type level, *capa-1(ok3065);nmur-1(ok1387)* double mutants exhibited survival comparable to that of *nmur-1(ok1387)* animals (Fig. 7C). This indicates that lacking CAPA-1 has no influence on *C. elegans* defense against *S. enterica*, and that CAPA-1 might not be involved in NMUR-1-mediated innate immune response to this pathogen. To further test this notion, we exposed wild-type and mutant animals to *S. enterica* on NGM media containing 0 or 1 µg/mL synthetic CAPA-1 peptide and scored their survival over time (Fig. 7D). In a parallel experiment, we first soaked wild-type and mutant animals in M9 buffer with 0 or 1 µg/mL synthetic CAPA-1 peptide for 1 hour, exposed the animals to *S. enterica* on NGM plates contain 0 or 1 µg/mL CAPA-1 peptide, and then scored for survival over time (Fig. S4). The results from both experiments showed that exogenous CAPA-1 peptide failed to influence the survival phenotype of *capa-1(ok3065)*, *nmur-1(ok1387)*, or *capa-1(ok3065);nmur-1(ok1387)* double mutant animals (Fig. 7D and S4), confirming that CAPA-1 is neither a ligand of NMUR-1 nor required in *C. elegans* defense against *S. enterica*. Although CAPA-1 is the only NMUR-1 ligand identified so far (Lindemans *et al*., 2009; Watteyne *et al*., 2020), these results suggest that other ligands may exist, and that NMUR-1 could bind to different ligands to regulate immune responses to distinct pathogens.

## Discussion

In the current study, we have shown that *C. elegans* lacking the neuronal GPCR NMUR-1 exhibited altered survival against various bacterial pathogens. By focusing on two pathogens, *E. faecalis* and *S. enterica*, as infecting agents, we have demonstrated that the altered survival was not caused by pathogen avoidance behavior but by differences in intestinal accumulation of pathogens, likely due to changes in innate immunity. Our RNA-seq analyses showed that functional loss of NMUR-1 negatively affects the expression of transcription factors involved in binding to RNA polymerase II regulatory regions, which, in turn, impacts transcriptional activity in *C. elegans*, as demonstrated by both *in vitro* and *in vivo* transcription assays. We also found that NMUR-1 regulates the expression of different sets of genes in *C. elegans* in response to *E. faecalis* and *S. enterica* exposure. In particular, C-type lectin genes were downregulated in *nmur-1(ok1387)* animals relative to wild-type animals exposed to *E. faecalis*, whereas UPR genes were upregulated following exposure to *S. enterica*. These genes have been previously implicated in conserved immune signaling pathways in response to pathogen infection (Sun *et al*., 2016). Functional assays with selected genes revealed that at least three C-type lectin genes (*clec-94*, *clec-208*, and *clec-263*) and two UPR genes (*T20D4.3* and *npl-4.2*) are important for *C. elegans* defense against *E. faecalis* and *S. enterica*, respectively. Taken together, we have demonstrated that NMUR-1 modulates *C. elegans* transcriptional activity by regulating the expression of transcription factors, which, in turn, controls the expression of distinct immune genes in response to different pathogens. These results uncovered a molecular basis for the specificity of *C. elegans* innate immunity. Given the evolutionary conservation of NMU/NMUR-1 signaling in immune regulation across multicellular organisms (Ye *et al*., 2021), our work could provide mechanistic insights into understanding the specificity of innate immunity in other animals, including mammals.

In vertebrates, a diverse array of PRRs that recognize various PAMPs have been implicated in the specificity of the innate immune system (Tan *et al*., 2014). Although *C. elegans* does not possess the majority of known PRRs, it can differentiate between different pathogens and launch distinct innate immune responses to specific pathogens (Irazoqui *et al*., 2010; Wong *et al*., 2007; Zarate-Potes *et al*., 2020). Recent behavior research in *C. elegans* indicates that its nervous system might play a major role in pathogen detection (Kim and Flavell, 2020). The nematode nervous system contains 32 chemosensory neurons (out of a total of 302 neurons in hermaphrodites), many of which have cilia exposed to the external environment that can directly sense pathogen-derived molecules in the environment (Bargmann, 2006). Moreover, *C. elegans* expresses approximately 1,300 GPCRs that may function as chemoreceptors in sensory neurons to detect pathogen cues, thus contributing to the specificity of innate immunity (Liu and Sun, 2021). In this context, a relevant question would be whether neuronal NMUR-1 mediates the specificity of innate immunity by directly interacting with bacterial molecules. We have shown that NMUR-1 has diverse effects on *C. elegans* defense against various bacterial pathogens (Fig. 1 and Fig. S1). If the hypothesis that NMUR-1 directly interacts with pathogens is correct, then this GPCR should be able to bind to various pathogen-derived molecules. Such scenario seems unlikely given that PRR-PAMP interactions follow a simple key-lock system, i.e., each receptor matches one pathogen-associated molecule or molecular pattern (Schulenburg *et al*., 2007). Therefore, it is more likely that NMUR-1 mediates the ability to differentiate between pathogens by controlling neural signaling than by recognizing bacterial molecules directly. Indeed, we have found that NMUR-1 regulates C-type lectin genes in response to *E. faecalis* infection but UPR genes in response to *S. enterica* infection; both of these genes are known to function in innate immune signaling pathways that fight infection (Sun *et al*., 2016).

How does NMUR-1, a single type of neuronal GPCR, distinguish different pathogens so that it can respond to some attacks with suppression of the innate immune response and amplify the response to others? A basic principle in neuroscience is that the interaction of multiple neural circuits mediates the functional output of a neural network and determines the final state of organ function (Tracey, 2014). Accordingly, the net effect of neural control on the immune response depends on the total sum of the interactions resulting from ligand-receptor signal transduction in a specific path. NMUR-1 is normally expressed in the spermathecae of the somatic gonad and several different types of sensory neurons as well as in interneurons (Maier *et al*., 2010). By using a fluorescent reporter transgene, we observed that distinct subsets of NMUR-1-expressing neurons are activated upon exposure to *E. faecalis* and *S. enterica* (data not shown), indicating that different neural circuits are involved in regulating the immune responses to these two pathogens. This notion is further supported by the fact that CAPA-1, the only NMUR-1 ligand identified so far, is required for the nematode immune response to *E. faecalis* but is dispensable for its response to *S. enterica* (Fig. 7). Future studies to decipher these different neuro-immune regulatory circuits would help us understand how signals pass through different components of the circuits to exert a net effect on immunity. An associative learning circuit mediated by CAPA-1/NMUR-1 signaling was recently uncovered in *C. elegans* in which sensory ASG neurons release CAPA-1, which signals via NMUR-1 in AFD sensory neurons to specifically mediate the retrieval of learned salt avoidance (Watteyne *et al*., 2020). In mice, cholinergic neurons in the intestine can sense and respond to infection by expressing NMU, which directly activates type 2 innate lymphoid cells in the gastrointestinal tract through NMUR1 to drive anti-parasitic immunity (Cardoso *et al*., 2017; Klose *et al*., 2017; Wallrapp *et al*., 2017). These studies provide useful information for deciphering NMUR-1-mediated neuro-immune regulatory circuits in *C. elegans*, which will be critical for further understanding how the nervous system regulates innate immunity to generate specificity by either suppressing or amplifying immune responses to different pathogens.

## Acknowledgements

We thank Dr. Danielle Garsin at the University of Texas Health Science Center at Houston for providing us with the *E. faecalis OG1RF::gfp* strain. We thank Dr. Joy Alcedo at Wayne State University for constructive discussions and for providing us with wild-type *N2* and *nmur-1(ok1387)* worm strains. Some worm strains were provided by the *Caenorhabditis* Genetics Center, which is funded by the NIH Office of Research Infrastructure Programs (P40 OD010440). This work was supported by NIH (R35GM124678 to J.S.), Department of Translational Medicine and Physiology, Elson S. Floyd College of Medicine, WSU-Spokane (Startup to Y.L.), and Research Foundation Flanders (G093419N to I.B.). The funders had no role in study design, data collection and interpretation, or the decision to submit the work for publication.

## Author contributions

P.W., S.W., J.W., C.C., D.S., Y.L., and I.B. designed and performed experiments and analyzed data. J.S. designed experiments and analyzed data. Y.L., J.S., and P.W. wrote the paper.

## Data and materials availability

The RNA-seq data have been deposited in NCBI’s SRA database through the GEO. The processed gene quantification files and differential expression files have been deposited in the GEO. All of the data can be accessed through the GEO with the accession number GSE154324. (Accession is currently private. To view the data, please go to https://www.ncbi.nlm.nih.gov/geo/query/acc.cgi?acc=GSE154324 Enter token ytghogwcpvwvbkv into the box). The *C. elegans* strains and plasmids constructed by us and primer sequences used for their construction are available upon request.

## Methods

### Nematode strains

The following *C. elegans* strains were maintained as hermaphrodites at 20°C, grown on modified NGM (0.35% instead of 0.25% peptone) with 30 units/mL nystatin and 15µg/mL tetracycline, and fed *E. coli* HT115. The wild-type animal strain was *C. elegans* Bristol N2. The *nmur-1(ok1387)*, *capa-1(ok3065)*, *CL2070(dvls70)*, and *TU3401* strains were obtained from the *Caenorhabditis elegans* Genetics Center (University of Minnesota, Minneapolis, MN). Some wild-type N2 and *nmur-1(ok1387)* used in this study were gifts from Dr. Joy Alcedo (Wayne State University, Detroit, MI). The *capa-1(ok3065);nmur-1(ok1387)*, *nmur-1(ok1387);CL2070(dvls70)*, and *nmur-1(ok1387);TU3401* mutant strains were constructed using standard genetic techniques.

### Bacterial strains

The following bacterial strains were grown using standard conditions (Lagier et al., 2015): *Escherichia coli* HT115, *Escherichia coli* OP50, *Salmonella enterica* strain SL1344, *S. enterica SL1344::gfp*, *Pseudomonas aeruginosa* strain PA14, *Yersinia pestis* strain KIM5, *Enterococcus faecalis* strain OG1RF, *E. faecalis OG1RF::gfp*, *Microbacterium nematophilum* strain CBX102, and *Staphylococcus aureus* strain NCTC8325.

### Survival assay

Wild-type and mutant animals were synchronized by egg-laying. Briefly, well-fed gravid adult animals were transferred to fresh *E. coli* HT115 seeded NGM plates and incubated for 45 minutes at 25°C. Adult animals were removed after 45 minutes, and the synchronized offspring were grown at 20°C for 65 hours. Bacterial lawns for the survival assays were prepared by placing a 30µL drop of an overnight culture of pathogenic bacteria on 3.5cm plates with either modified NGM or brain heart infusion (BHI) media for *E. faecalis* OG1RF. Full-lawn survival assays were prepared by spreading 30µL of overnight bacterial culture over the complete surface of each 3.5cm plate. Plates were incubated at 37°C for 15∼16 hours, cooled to room temperature, and then seeded with synchronized 65-hour-old young adult animals. The survival assays were performed at 25°C, and live animals were transferred daily to fresh plates until egg laying ceased. Animals were scored at the times indicated and were considered dead when they failed to respond to touch.

### Lawn occupancy assay

Synchronized 65-hour-old animals and bacteria plates were prepared using the method described above. Synchronized animals were placed in the center of the bacterial lawn and allows to crawl freely on the plate. The number of animals on the bacteria lawn were counted at 4, 8, 12, 24, and 36 hours post-exposure. The assays were performed at 25°C and dead animals were censored. The results were calculated as a percentage of animals on the bacteria lawn versus the total number of animals on the plates at the given time point.

### Profiling bacterial accumulation in the nematode intestine

Synchronized 65-hour-old animals were prepared using the method describe above. GFP-tagged bacteria were prepared using the method described above with the addition of antibiotics in the media plates. Synchronized animals were transferred to the plates seeded with GFP-tagged bacteria and incubated for 24 hours at 25°C. Following incubation, the animals were transferred to an empty NGM plate for 15 minutes to eliminate any fluorescent bacteria stuck to the body. The animals were then transferred to a new, empty NGM plate and into a droplet of M9 buffer to further wash away any bacteria remaining on the body. Animals were anesthetized using 25mM Sodium Azide and visualized using a Zeiss Axio Imager M2 stereoscopic microscope.

### Quantification of intestinal colonization

Synchronized 65-hour-old animals were prepared using the method describe above. GFP-tagged bacteria were prepared using the method described above with the addition of antibiotics in the media plates. Synchronized animals were transferred to the plates seeded with GFP-tagged bacteria and incubated for 24 hours at 25°C. Ten animals were transferred to a 1.5mL tube containing a solution of 25mM Sodium Azide with 1mg/mL ampicillin and kanamycin. The animals were then soaked for 45 minutes to remove any external bacteria. After 45 minutes, the animals were washed three times with PBS containing 0.1% Triton to remove the remaining antibiotics, and ground in 100µL of PBS with 0.1% Triton. Serial dilutions of the lysates (10^-1^, 10^-2^, 10^-3^, 10^-4^, and 10^-5^) were plated with rifampicin to select for GFP-tagged bacteria and the plates were incubated at 37°C for 24 hours.

### Profiling pumping and defecation rates

Synchronized 65-hour-old animals, and bacteria plates were prepared using the method described above. Synchronized animals were transferred to plates seeded with pathogenic bacteria and incubated for 24 hours at 25°C. Pumping rates were counted by observing the contraction/relaxation cycles of the terminal pharyngeal bulb using a stereoscopic microscope. Cycles were counted for 30 second durations, three times for ten individual animals per condition. Defecation rates were measured by timing the span between the contraction of the rectal muscles and subsequent excretion using a stereoscopic microscope. Cycles were counted five times for ten individual animals per condition.

### RNA interference

RNAi was conducted by feeding synchronized L3 larval *C. elegans E. coli* strain HT115(DE3) expressing double-stranded RNA (dsRNA) that was homologous to a target gene (Fraser et al., 2000; Timmons and Fire, 1998). *E. coli* with the appropriate dsRNA vector were grown in LB broth containing ampicillin (100µg/mL) at 37°C for 15∼16 hours and plated on modified NGM plates containing 100µg/mL ampicillin and 3mM isopropyl β-D-thiogalactoside (IPTG). The bacteria were allowed to grow for 15∼16 hours at 37°C. The plates were cooled away from direct light before the synchronized L3 larval animals were placed on the bacteria. The animals were incubated at 20°C for 24 hours or until the animals were 65 hours old. *unc-22* RNAi was included as a positive control in all experiments to account for RNAi efficiency.

### Synthetic NMUR-1 ligand administration

Synchronized 65-hour-old animals, and bacteria plates were prepared using the method described above. Synchronized animals were transferred to modified NGM or BHI bacteria plates containing 0, 1, or 5µg/mL oligopeptide AFFYTPRI-NH2. The survival assays were performed at 25°C and live animals were transferred daily to fresh plates until egg laying ceased. The animals were scored at the times indicated and were considered dead when they failed to respond to touch.

Wild-type and mutant animals were also lysed using a solution of sodium hydroxide and bleach (ratio 5:2), washed, and eggs were synchronized for 22 hours in S-basal liquid medium at room temperature. Synchronized L1 larval animals were transferred onto modified NGM plates seeded with *E. coli* HT115 and grown at 20°C for 65 hours. Synchronized animals were washed in M9 buffer and soaked for one hour in M9 buffer containing 0, 1, or 5µg/mL oligopeptide AFFYTPRI-NH2. Soaked animals were transferred to modified NGM medium containing 0, 1, or 5µg/mL oligopeptide AFFYTPRI-NH2 seeded with 30µL of fresh pathogen culture that was incubated at 37°C for 15∼16 hours. The survival assays were performed at 25°C and live animals were transferred daily to fresh plates until egg laying ceased. The animals were scored at the times indicated and were considered dead when they failed to respond to touch.

### RNA isolation

Gravid adult wild-type and *nmur-1(ok1387)* animals were lysed using a solution of sodium hydroxide and bleach (ratio 5:2), washed, and eggs were synchronized for 22 hours in S-basal liquid medium at room temperature. Synchronized L1 larval animals were transferred onto modified NGM plates seeded with *E. coli* HT115 and grown at 20°C for 48 hours until the animals had reached the L4 larval stage. The animals were collected and washed with M9 buffer before being transferred to plates containing *E. coli* HT115 or pathogenic bacteria for 24 hours at 25°C. After 24 hours, animals were collected and washed with M9 buffer, and RNA was extracted using QIAzol (Qiagen). Total RNA was isolated using the RNeasy mini kit (Qiagen) following the protocol provided by the manufacturer.

### Quantitative real-time PCR (qRT-PCR)

Total RNA was obtained as described above. 2µg of RNA were used to generate cDNA using the Applied Biosystems High-Capacity cDNA Reverse Transcription Kit. qRT-PCR was conducted by following the prescribed protocol for PowerUp SYBR Green (Applied Biosystems) on an Applied Biosystems StepOnePlus real-time PCR machine. 10µL reactions were set up following the manufacturer’s recommendations, and 20ng of cDNA were used per reaction. Relative fold-changes for transcripts were calculated using the comparative *C_T_*(2*^-ΔΔCT^*) method and were normalized to pan-actin (*act-1, −3, −4*). Amplification cycle thresholds were determined by the StepOnePlus software. All samples were run in triplicate. Primer sequences are available upon request.

### RNA sequencing

Total RNA samples were obtained as described above and was submitted to the WSU Genomics Core for RNA-seq analyses. RNA-seq and related bioinformatic analyses were done following our published protocol (Sellegounder *et al*., 2019). The raw sequence data (FASTQ files) were deposited in NCBI’s SRA database through the GEO. The processed gene quantification files and differential expression files were deposited in GEO. All of these data can be accessed through GEO with the accession number GSE154324.

### In vitro transcription assay

In vitro transcription assays were performed following our published protocol (Wibisono *et al*., 2020). Briefly, nuclear extracts were prepared from synchronized young adult animals using Balch homogenization. The nuclear extracts were then incubated with PESDNA (a linear DNA template containing the worm *Δpes-10* promoter) and other transcription components at 30°C for 30 minutes, followed by RNA purification and reverse transcription. The resulting cDNA was either amplified by PCR and detected by gel electrophoresis or quantified by qRT-PCR using the PowerUp SYBR green qPCR kit (Applied Biosystems, catalog # A25918).

### Plasmid construction and transgenic animal generation

A *clec-94* genomic DNA fragment (4,247 bp) was amplified by PCR from mixed stage wild-type *C. elegans* using the following primers: 5’-ATTGTCGACGTACAGTCCCACTACTTGTTGC - 3’ and 5’-CTAGGATCCCTAACATTGGCCGGGAAAGAG −3’. PCR products were then cloned into the *pPD95.77* vector (Fire Lab *C. elegans* vector kit; Addgene, Cambridge, MA) via the *SalI* and *BamHI* sites. The resulting plasmid pPW01 (*clec-94p::clec-94::SL2::gfp*) was microinjected into wild-type animals at 10ng/µL to generate strain JRS60 (Table S3). The *clec-94* genomic rescue strain JRS63 (Table S3) was generated by crossing JRS60 with *nmur-1(ok1387)* animals.

A *clec-208* genomic DNA fragment (3,556 bp) was amplified by PCR from mixed stage wild-type *C. elegans* using the following primers: 5’-TTTGTCGACTTTCTGTTCTTGCTACTCTCTACC −3’ and 5’-TTTTCTAGACTGGCTCGTTCTTAGAGACC −3’. PCR products were then cloned into the *pPD95.77* vector via the *SalI* and *XbaI* sites. The resulting plasmid, pPW02 (*clec-208p::clec-208::SL2::gfp*), was microinjected into wild-type animals at 50ng/µL with *unc-122p::mCherry* at 20ng/µl as a co-injection marker to generate strain JRS61 (Table S3). The *clec-208* genomic rescue strain JRS64 (Table S3) was generated by crossing JRS61 with *nmur-1(ok1387)* animals.

A *clec-263* genomic DNA fragment (5,389 bp) was amplified by PCR from mixed stage wild-type *C. elegans* using the following primers: 5’-TTTGTCGACGCGATGGGTGGTTGTATTATTC −3’ and 5’-CAAGGATCCAAATCGTATTTCCGTCGTTGGTC −3’. PCR products were then cloned into the *pPD95.77* vector via the *SalI* and *BamHI* sites. The resulting plasmid, pPW03 (*clec-263p::clec-263::SL2::gfp*), was microinjected into wild-type animals at 30ng/µL with *unc-122p::mCherry* at 20ng/µl as a co-injection marker to generate strain JRS62 (Table S3). The *clec-263* genomic rescue strain JRS65 (Table S3) was generated by crossing JRS62 with *nmur-1(ok1387)* animals.

A *T20D4.3* genomic DNA fragment (7,665 bp) was amplified by PCR from mixed stage wild-type *C. elegans* using oligonucleotides 5’-GTACTGCAGCTGTGTCGCCAGAATGATTG −3’ and 5’-GATTGGCCATGTGAATGTTAAGAAGGCGTG −3’. PCR products were then cloned into vector pPD95.77 (Fire Lab C. elegans vector kit; Addgene, Cambridge, MA) via the *SalI* and *BamHI* sites. The resulting plasmid, pPW04 (*T20D4.3p::T20D4.3::SL2::gfp*), was microinjected into wild-type animals at 50ng/µL with *myo-3p::mCherry* at 10ng/µl as a co-injection marker to generate overexpression strain JRS78 (Table S3). The *T20D4.3* overexpression strain JRS79 (Table S3) was generated by crossing JRS78 with *nmur-1(ok1387)*.

### Statistical analysis

Survival curves were plotted using GraphPad PRISM (version 9) computer software. Survival was considered different from the appropriate control indicated in the main text when *P* <0.05. PRISM uses the product limit or Kaplan-Meier method to calculate survival fractions and the log-rank test, which is equivalent to the Mantel-Haenszel test, to compare survival curves. Occupancy assays, CFU assays, and qRT-PCR results were analyzed using two-sample *t*-tests for independent samples; *P-*values <0.05 are considered significant. All experiments were repeated at least three times, unless otherwise indicated.

## Supplemental information

**Supplemental materials include Figs. S1 to S4 and Tables S1 to S3**

**Fig. S1. Functional loss of NMUR-1 differentially affects *C. elegans* survival against various bacteria.** WT and *nmur-1(ok1387)* animals were exposed to *E. coli* OP50 **(A)**, *P. aeruginosa* PA14 **(B)**, *Y. pestis* KIM5 **(C)**, *M. nematophilum* CBX102 **(D)**, or *S. aureus* NCTC8325 **(E)** and scored for survival over time. Each graph is a combination of three independent experiments. Each experiment included *N* = 60 animals per strain. *P*-values represent the significance level of mutant survival relative to WT, *P* = 0.0011 in (A), *P* = 0.8084 in (B), *P* < 0.0001 in (C), *P* = 0.2401 in (D), and *P* = 0.0678 in (E).

**Fig. S2. Functional loss of NMUR-1 does not affect *C. elegans* survival against heat-killed *E. faecalis* or heat-killed *S. enterica*. WT and *nmur-1(ok1387)* animals were exposed to heat-killed *E. faecalis* (A)** or heat-killed *S. enterica* **(B)** and scored for survival over time. Each graph is a combination of three independent experiments. Each experiment included *N* = 60 animals per strain. *P*-value represents the significance level of the survival of mutants relative to WT, *P* = 0.2191 in (A) and *P* = 0.2121 in (B).

**Fig. S3. Functional loss of NMUR-1 differentially affects *C. elegans* survival against *E. faecalis* and *S. enterica* in full-lawn assays. WT and *nmur-1(ok1387)* animals were exposed to a full lawn of *E. faecalis* (A)** or *S. enterica* **(B)** and scored for survival over time. Each graph is a combination of three independent experiments. Each experiment included *N* = 60 animals per strain. *P*-values represent the significance level of the survival of mutants relative to WT, *P* < 0.0001 in (A) and *P* < 0.0001 in (B).

**Fig. S4. Exogenous CAPA-1 peptide does not alter the survival phenotype of *capa-1(ok3065)* mutants against *S. enterica*.** WT, *nmur-1(ok1387), capa-1(ok3065),* and *capa-1(ok3065);nmur-1(ok1387)* animals were soaked in M9 buffer with either 0 or 1µg/mL synthetic CAPA-1 peptide for one hour, exposed to *S. enterica* on plates contain either 0 or 1µg/mL synthetic CAPA-1 peptide, and scored for survival over time. The graph is a combination of three independent experiments. Each experiment included *N =* 60 animals per strain. *P*-values represent the significance level of the mutant survival relative to the WT+1µg/mL: *nmur-1(ok1387)+*1µg/mL*, P* < 0.0001; *capa-1(ok3065) +*1µg/mL, *P* = 0.3509; *capa-1(ok3065);nmur-1(ok1387)+*1µg/mL, *P* < 0.0001.

**Table S1. Enrichment of molecular functions revealed by GO analysis of upregulated genes in *nmur-1(ok1387)* animals relative to wild-type animals.**

**Table S2. Enrichment of biological processes revealed by GO analysis of upregulated genes in nmur-1(ok1387) animals relative to wild-type animals exposed to E. faecalis.**

**Table S3. List of transgenic *C. elegans* strains generated in this study.**

