## Supplemental information for "Neuronal GPCR regulates the specificity of innate immunity against pathogen infection"

**Fig. S1**

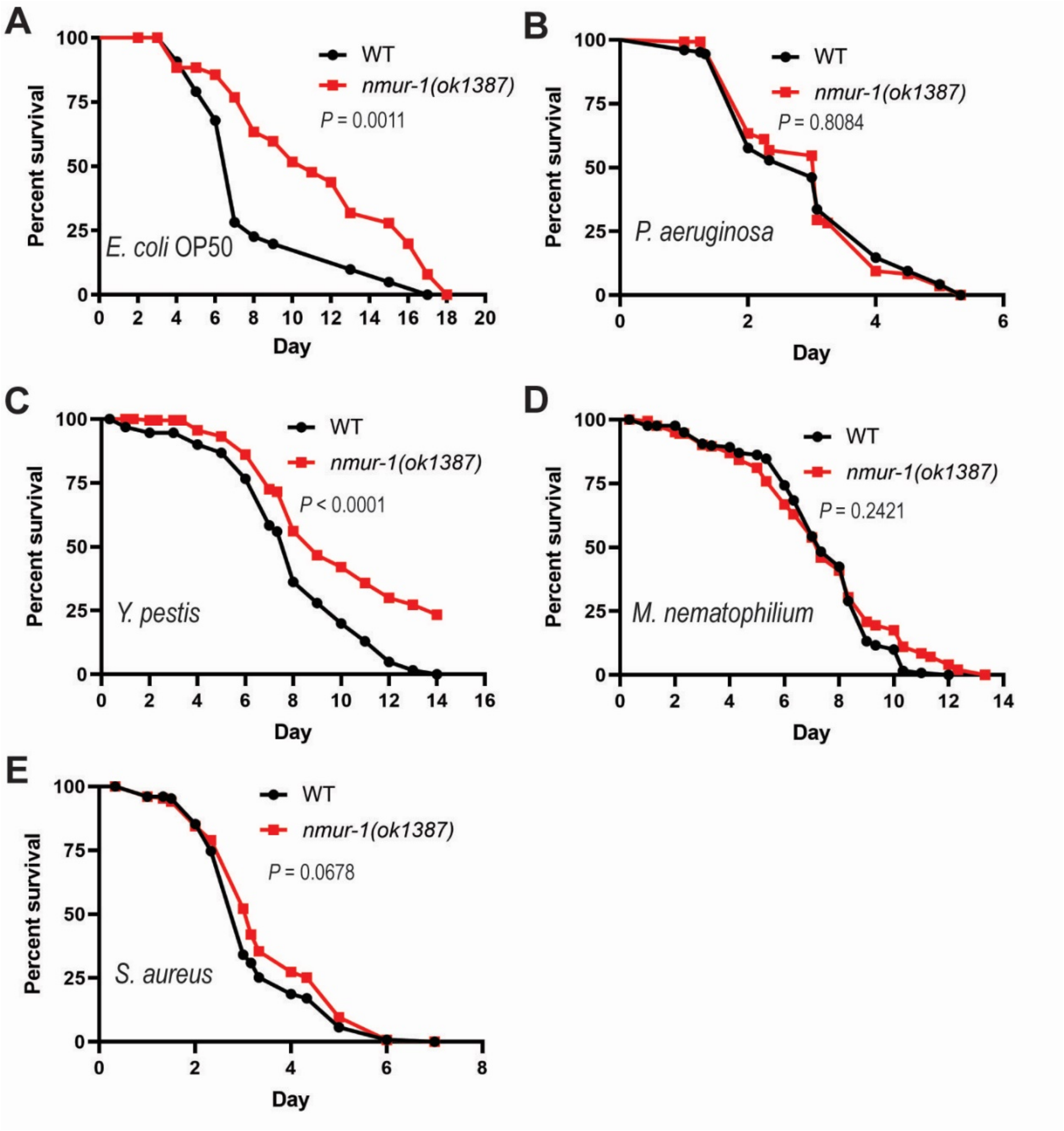


**Fig. S1. Functional loss of NMUR-1 differentially affects *C. elegans* survival against various bacteria.** WT and *nmur-1(ok1387)* animals were exposed to *E. coli* OP50 **(A)**, *P. aeruginosa* PA14 **(B)**, *Y. pestis* KIM5 **(C)**, *M. nematophilum* CBX102 **(D)**, or *S. aureus* NCTC8325 **(E)** and scored for survival over time. Each graph is a combination of three independent experiments. Each experiment included $N = 60$ animals per strain. *P*-values represent the significance level of mutant survival relative to WT, $P = 0.0011$ in (A), $P = 0.8084$ in (B), $P < 0.0001$ in (C), $P = 0.2401$ in (D), and $P = 0.0678$ in (E).

**Fig. S2**

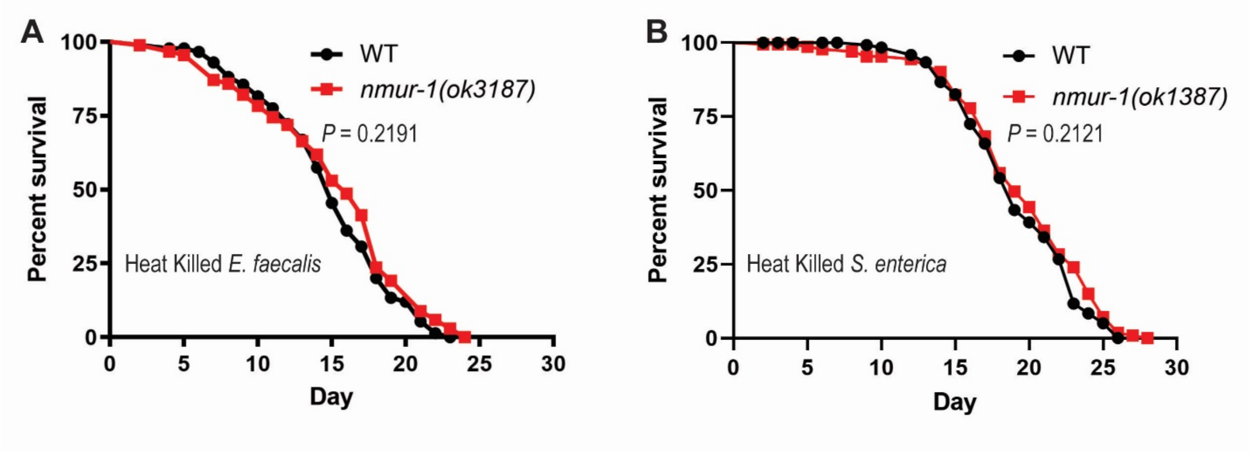


**Fig. S2. Functional loss of NMUR-1 does not affect *C. elegans* survival against heat-killed *E. faecalis* or heat-killed *S. enterica*.** WT and *nmur-1(ok1387)* animals were exposed to heat-killed *E. faecalis* **(A)** or heat-killed *S. enterica* **(B)** and scored for survival over time. Each graph is a combination of three independent experiments. Each experiment included $N = 60$ animals per strain. *P*-value represents the significance level of the survival of mutants relative to WT, $P = 0.2191$ in (A) and $P = 0.2121$ in (B).

**Fig. S3**

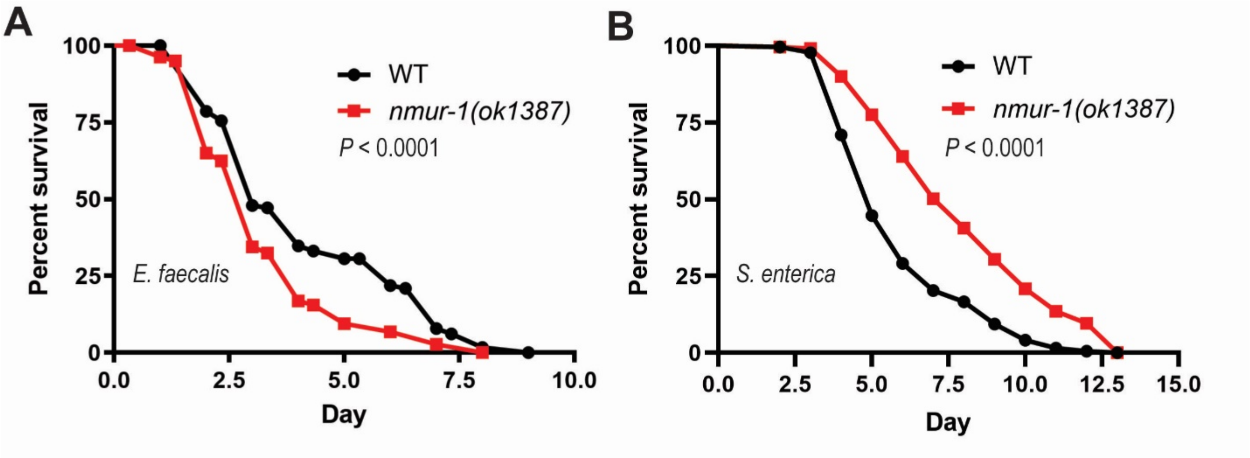


**Fig. S3. Functional loss of NMUR-1 differentially affects *C. elegans* survival against *E. faecalis* and *S. enterica* in full-lawn assays.** WT and *nmur-1(ok1387)* animals were exposed to a full lawn of *E. faecalis* **(A)** or *S. enterica* **(B)** and scored for survival over time. Each graph is a combination of three independent experiments. Each experiment included $N = 60$ animals per strain. *P*-values represent the significance level of the survival of mutants relative to WT, $P < 0.0001$ in (A) and $P < 0.0001$ in (B).

**Fig. S4**

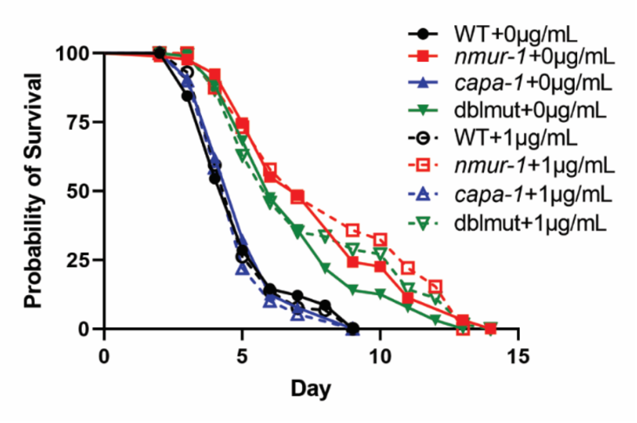


**Fig. S4. Exogenous CAPA-1 peptide does not alter the survival phenotype of *capa-1(ok3065)* mutants on *S. enterica*.** WT, *nmur-1(ok1387), capa-1(ok3065),* and *capa-1(ok3065);nmur-1(ok1387)* animals were soaked in M9 buffer with either 0 or 1µg/mL synthetic CAPA-1 peptide for one hour and exposed to *S. enterica* on plates contain either 0 or 1µg/mL synthetic CAPA-1 peptide and scored for survival over time. The graph is a combination of three independent experiments. Each experiment included $N = 60$ animals per strain. $P$-values represent the significance level of the mutant survival relative to the WT+1µg/mL: *nmur-1(ok1387)*+1µg/mL, $P < 0.0001$; *capa-1(ok3065)* +1µg/mL, $P = 0.3509$; *capa-1(ok3065);nmur-1(ok1387)*+1µg/mL, $P < 0.0001$.

**Table S1. Enrichment of molecular functions revealed by GO analysis of upregulated genes in *nmur-1(ok1387)* animals relative to wild-type animals.**

| GO term | Description | P-value[a] | FDR q-value[b] | Enrichment (N, B, n, b)[c] |
|---|---|---|---|---|
| GO:0004721 | phosphoprotein phosphatase activity | 1.21E-40 | 3.08E-37 | 5.37 (9948,161,921,80) |
| GO:0004725 | protein tyrosine phosphatase activity | 6.28E-37 | 7.97E-34 | 6.95 (9948,87,921,56) |
| GO:0042302 | structural constituent of cuticle | 1.05E-34 | 8.86E-32 | 5.62 (9948,125,921,65) |
| GO:0140096 | catalytic activity, acting on a protein | 1.97E-33 | 1.25E-30 | 2.17 (9948,1133,921,228) |
| GO:0004672 | protein kinase activity | 2.29E-30 | 1.16E-27 | 3.19 (9948,376,921,111) |
| GO:0016791 | phosphatase activity | 4.24E-30 | 1.79E-27 | 4.08 (9948,212,921,80) |
| GO:0042578 | phosphoric ester hydrolase activity | 3.37E-25 | 1.22E-22 | 3.53 (9948,245,921,80) |
| GO:0016773 | phosphotransferase activity, alcohol group as acceptor | 9.08E-25 | 2.88E-22 | 2.78 (9948,431,921,111) |
| GO:0004715 | non-membrane spanning protein tyrosine kinase activity | 2.07E-21 | 5.83E-19 | 7.36 (9948,44,921,30) |
| GO:0004674 | protein serine/threonine kinase activity | 2.35E-21 | 5.96E-19 | 3.12 (9948,277,921,80) |
| GO:0016301 | kinase activity | 2.42E-21 | 5.59E-19 | 2.52 (9948,485,921,113) |
| GO:0016772 | transferase activity, transferring phosphorus-containing groups | 1.34E-15 | 2.84E-13 | 2.13 (9948,579,921,114) |
| GO:0016788 | hydrolase activity, acting on ester bonds | 1.19E-14 | 2.33E-12 | 2.22 (9948,471,921,97) |
| GO:0004713 | protein tyrosine kinase activity | 9.57E-13 | 1.74E-10 | 4.19 (9948,80,921,31) |
| GO:0004722 | protein serine/threonine phosphatase activity | 2.86E-11 | 4.84E-09 | 4.63 (9948,56,921,24) |
| GO:0005198 | structural molecule activity | 4.50E-09 | 7.14E-07 | 2.08 (9948,342,921,66) |
| GO:0005524 | ATP binding | 5.16E-08 | 7.71E-06 | 1.61 (9948,798,921,119) |
| GO:0032559 | adenyl ribonucleotide binding | 1.31E-07 | 1.85E-05 | 1.58 (9948,820,921,120) |
| GO:0030554 | adenyl nucleotide binding | 1.60E-07 | 2.14E-05 | 1.57 (9948,823,921,120) |
| GO:0008144 | drug binding | 1.52E-06 | 1.93E-04 | 1.49 (9948,912,921,126) |
| GO:0035639 | purine ribonucleoside triphosphate binding | 7.87E-05 | 9.51E-03 | 1.38 (9948,960,921,123) |
| GO:0032555 | purine ribonucleotide binding | 1.46E-04 | 1.68E-02 | 1.36 (9948,983,921,124) |
| GO:0017076 | purine nucleotide binding | 1.74E-04 | 1.92E-02 | 1.36 (9948,987,921,124) |
| GO:0005102 | signaling receptor binding | 1.89E-04 | 1.99E-02 | 1.92 (9948,186,921,33) |
| GO:0032553 | ribonucleotide binding | 2.89E-04 | 2.94E-02 | 1.34 (9948,999,921,124) |
| GO:0042329 | structural constituent of collagen and cuticulin-based cuticle | 3.51E-04 | 3.43E-02 | 5.40 (9948,12,921,6) |
| GO:0030246 | carbohydrate binding | 4.64E-04 | 4.37E-02 | 1.94 (9948,156,921,28) |

[a]P-value is computed according to the mHG model (Eden *et al.* 2007 PLoS Comp Bio 3(3):e39).
[b]FDR q-value is the correction of the above p-value for multiple testing using the Benjamini and Hochberg method (Benjamini and Hochberg 1995 J R Statist Soc B 57(1):289-300).
[c]Enrichment (N, B, n, b) is defined as follows:
N - total number of genes
B - total number of genes associated with a specific GO term
n - number of genes in the target set
b - number of genes in the intersection
Enrichment = (b/n) / (B/N)

**Table S2. Enrichment of biological processes revealed by GO analysis of upregulated genes in *nmur-1(ok1387)* animals relative to wild-type animals exposed to *E. faecalis*.**

| GO term | Description | P-value[a] | FDR q-value[b] | Enrichment (N, B, n, b)[c] |
|---|---|---|---|---|
| GO:0050830 | defense response to Gram-positive bacterium | 1.24E-11 | 7.49E-08 | 15.32 (10357,61,133,12) |
| GO:0042742 | defense response to bacterium | 3.97E-09 | 1.20E-05 | 7.57 (10357,144,133,14) |
| GO:0009617 | response to bacterium | 3.97E-09 | 7.98E-06 | 7.57 (10357,144,133,14) |
| GO:0098542 | defense response to other organism | 2.70E-08 | 4.07E-05 | 6.53 (10357,167,133,14) |
| GO:0043207 | response to external biotic stimulus | 2.91E-08 | 3.52E-05 | 6.49 (10357,168,133,14) |
| GO:0009607 | response to biotic stimulus | 2.91E-08 | 2.93E-05 | 6.49 (10357,168,133,14) |
| GO:0051707 | response to other organism | 2.91E-08 | 2.51E-05 | 6.49 (10357,168,133,14) |
| GO:0051704 | multi-organism process | 2.02E-07 | 1.52E-04 | 5.56 (10357,196,133,14) |
| GO:0009605 | response to external stimulus | 9.53E-06 | 6.39E-03 | 4.02 (10357,271,133,14) |
| GO:0042338 | cuticle development involved in collagen and cuticulin-based cuticle molting cycle | 1.13E-05 | 6.83E-03 | 16.22 (10357,24,133,5) |
| GO:0006952 | defense response | 3.61E-05 | 1.98E-02 | 3.38 (10357,346,133,15) |

[a]P-value is computed according to the mHG model (Eden *et al.* 2007 PLoS Comp Bio 3(3):e39).
[b]FDR q-value is the correction of the above p-value for multiple testing using the Benjamini and Hochberg method (Benjamini and Hochberg 1995 J R Statist Soc B 57(1):289-300).
[c]Enrichment (N, B, n, b) is defined as follows:
N - total number of genes
B - total number of genes associated with a specific GO term
n - number of genes in the target set
b - number of genes in the intersection
Enrichment = (b/n) / (B/N)

**Table S3. List of transgenic *C. elegans* strains generated in this study.**

| Strain | Plasmid and genetic background | Host | Note |
|---|---|---|---|
| JRS60 | pPW01 *clec-94p(4.2kb)::clec-94 gDNA::SL2::gfp* | *N2* | *clec-94* gene over-expression in *N2* animals |
| JRS61 | pPW02 *clec-208p(3.6kb)::clec-208 gDNA::SL2::gfp* | *N2* | *clec-208* gene over-expression in *N2* animals |
| JRS62 | pPW03 *clec-263p(5.4kb)::clec-263 gDNA::SL2::gfp* | *N2* | *clec-263* gene over-expression in *N2* animals |
| JRS63 | pPW01 *clec-94p(4.2kb)::clec-94 gDNA::SL2::gfp* | *nmur-1(ok1387)* | *clec-94* genomic rescue driven by *clec-94* promoter |
| JRS64 | pPW02 *clec-208p(3.6kb)::clec-208 gDNA::SL2::gfp* | *nmur-1(ok1387)* | *clec-208* genomic rescue driven by *clec-208* promoter |
| JRS65 | pPW03 *clec-263p(5.4kb)::clec-263 gDNA::SL2::gfp* | *nmur-1(ok1387)* | *clec-263* genomic rescue driven by *clec-263* promoter |
| JRS78 | pPW04 *T20D4.3p(7.7kb)::T20D4.3 gDNA::SL2::gfp* | *N2* | *T20D4.3* gene over-expression in *N2* animals |
| JRS79 | pPW04 *T20D4.3p(7.7kb)::T20D4.3 gDNA::SL2::gfp* | *nmur-1(ok1387)* | *T20D4.3* genomic rescue driven by *T20D4.3* promoter |
